# NAD^+^ limitation and α-ketoglutarate depletion define cancer induced muscle atrophy

**DOI:** 10.64898/2026.10.15.759464

**Authors:** Vassilis Papalazarou, Jack Coomes, Lucia Rossi, Valentina M. Lionello, Izadora Furlani, Francesco Saverio Tedesco, Karen H. Vousden

**Affiliations:** Department of Cell & Developmental Biology, Division of Biosciences, University College London, London WC1E 6DE, UK; Dubowitz Neuromuscular Centre, Great Ormond Street Hospital and UCL Great Ormond Street Institute of Child Health, London, UK; The Francis Crick Institute, 1 Midland Road, London NW1 1AT UK

**Keywords:** Muscle atrophy, pancreas cancer, cachexia, lactate, NAD+/NADH ratio, α-ketoglutarate

## Abstract

Tumour-host crosstalk is an emerging hallmark of malignancy. It is increasingly recognised that cancer alters non-tumour bearing distant organs, with important consequences for disease progression and survival. Skeletal muscle loss, a defining feature of cancer-associated cachexia, contributes to morbidity and reduced quality of life, yet its underlying mechanisms remain poorly understood. In this study we analysed the secretome of pancreatic cancer cells and identified a metabolite-driven signal that induces myotube atrophy. Our data suggests that atrophy, which seems to involve cytoskeletal stress and increased myosin contractility, arises from cancer cell-dependent reductive metabolic rewiring. Pancreatic cancer cells conditioned their environment with elevated levels of lactate, generating a lactate-rich extracellular milieu that decrease the NAD^+^/NADH ratio in myotubes. This redox imbalance impaired oxidative metabolism, decreasing α-ketoglutarate (αKG) levels and disrupted αKG-dependent processes. Restoration of NAD^+^ or αKG levels rescued myotube atrophy. Skeletal muscle from cancer-associated cachexia mouse models exhibited reduced αKG levels. Collectively, these findings identify a targetable metabolite-mediated mechanism of tumour-muscle crosstalk.

## INTRODUCTION

Cancer cachexia is a multifaceted systemic disorder that affects up to 80% of cancer patients and directly accounting for more than 20% of cancer related deaths^1^. In pancreatic cancer, the fourth leading cause of cancer-related mortality in Western countries, severe tissue wasting contributes for up to 80% of death in patients with advanced disease^2^. Skeletal muscle loss is central to the current definition of cancer-associated cachexia, described as ‘a multifactorial syndrome characterized by an ongoing loss of skeletal muscle mass (with or without loss of fat mass) that cannot be fully reversed by conventional nutritional support and leads to progressive functional impairment^3^. In pancreatic ductal adenocarcinoma (PDAC), evidence from both mouse models and patients indicates that adipose and muscle tissue wasting are early events^4,5^, while muscle loss is associated with poorer life expectancy^6^. Early tissue wasting has been associated with elevated circulating branched chain amino acids (BCAAs), likely derived from systemic protein breakdown^7^. However, it is important to note that PDAC mouse models predominantly exhibit adipose tissue loss, with comparatively modest muscle wasting^5^.

Loss of lean body mass arises from a heterogeneous palette of metabolic, hormonal, behavioural and inflammatory mediators^8^. These include pro-inflammatory cytokines, such as tumour necrosis factor-α (TNF-α)^9^, IL-6^10^, and Activin-A^11^, as well as factors acting on the central nervous system (CNS) (e.g. GDF15)^12^, and inter-organ communication signals, including adipose-muscle crosstalk^13^. Recent work suggests the final cachectic outcome is the result of complex interactions between the cancer, the central nervous system and various organs that lead to modulation in appetite in addition to direct tissue wasting^14^. Whether these systemic factors directly drive muscle specific effects or instead reflect advanced, irreversible disease remains unclear. Their mechanisms of action in muscle are also incompletely understood, with apoptosis, autophagy and proteolysis mechanisms all proposed to contribute^15–17^. There are currently no approved therapies for cachexia, however a humanized monoclonal antibody against GDF-15 improved weight and physical activity in cancer patients with cachexia and elevated circulating GDF-15 levels, in a phase 2 clinical trial^18^.

Muscle atrophy is generally considered to result from disrupted proteostasis, reflecting an imbalance between protein synthesis and degradation^16,19^. This is supported by the characterisation of ubiquitin-mediated degradation pathways and the induction of so-called atrogenes^20–22^. However, in the context of skeletal muscle repair and regeneration^19,23^, it remains unclear whether these processes represent terminal events in fibre degradation, compensatory clearance mechanisms, or primary drivers of muscle loss. In the process of cancer cachexia, tumor derived inflammatory molecules and cytokines have been implicated in the activation transcriptional responses that drive muscle proteolysis^24^, in addition to promoting mitochondrial dysfunction that seems to be a hallmark of cachectic muscle^25^.

Cachexia is increasingly recognised as a metabolic disease. Recent studies using multi-organ and multi-omics approaches have begun to delineate metabolic crosstalk between tumours and distant tissues^26–29^. Several metabolic alterations have been associated with cachexia, including certain changes in amino acids and lipid species^30,31^. A large scale screen of many of these metabolites showed that the oncometabolite D-hydroxygluterate – which accumulates in IDH mutant tumours – can promote muscle atrophy^32^. However, our understanding of the role of other metabolites in tumors lacking IDH mutations remains unclear. Given this limited mechanistic understanding, together with the lack of models that fully recapitulate all aspects of the disease, it is not surprising that effective therapies remain unavailable and cachexia is largely irreversible despite efforts for nutritional support.

To circumvent this complexity, we employed *in vitro* 2D and 3D mouse and human models to investigate how PDAC-derived factors influence muscle cells. We found that PDAC secreted metabolites, but not proteins or lipids, directly induce myotube atrophy. This effect was not primarily driven by increased proteolysis but was instead associated with mechanical stress and enhanced contractility. Among these metabolites, lactate was the most abundant factor in the PDAC secretome and a key driver of myotube atrophy. Specifically, increased PDAC-derived extracellular lactate impaired lactate flux in myotubes, leading to intracellular lactate accumulation and a reduction in the NAD^+^/NADH ratio. This shift limited α-ketoglutarate (αKG) synthesis, with downstream effects on αKG-dependent processes. Restoration of NAD^+^ or αKG levels rescued PDAC-induced atrophy. Consistently, skeletal muscle from cachectic mouse models exhibited a reduced αKG/succinate ratio.

Together, these findings provide a mechanistic framework for PDAC-muscle metabolic crosstalk and identify a targetable metabolic vulnerability underlying cancer-associated muscle loss.

## RESULTS

### Metabolites secreted by PDAC cells induce myotube atrophy

To model crosstalk between pancreatic cancer cells and myotubes *in vitro*, C2C12 progenitor cells were differentiated into myotubes and subsequently exposed to conditioned media derived from murine pancreatic cancer cell lines generated from genetically engineered models of pancreatic cancer harbouring Kras G12D mutation together with either p53 loss (‘KFC’) or TP53 R172H mutation (‘KPC’) (Extended Data Fig. 1A). In parallel, conditioned media from human pancreatic cancer cell lines carrying p53 mutations (PANC1:R273H, MiaPaCa2:R248W) or p53 loss (AsPC1) were used. To minimise differences in proliferation rates across cell lines and promote the accumulation of tumour-derived secreted factors, cells were cultured in low-serum conditions (1% dFBS) for 24 hours prior to media collection. Pre-conditioned media or media conditioned by undifferentiated C2C12 cells served as controls. For all experiments conditioned media was applied at a 50% dilution in myotube maintenance medium to prevent nutrient deprivation (Figure 1A).

**Figure 1:**
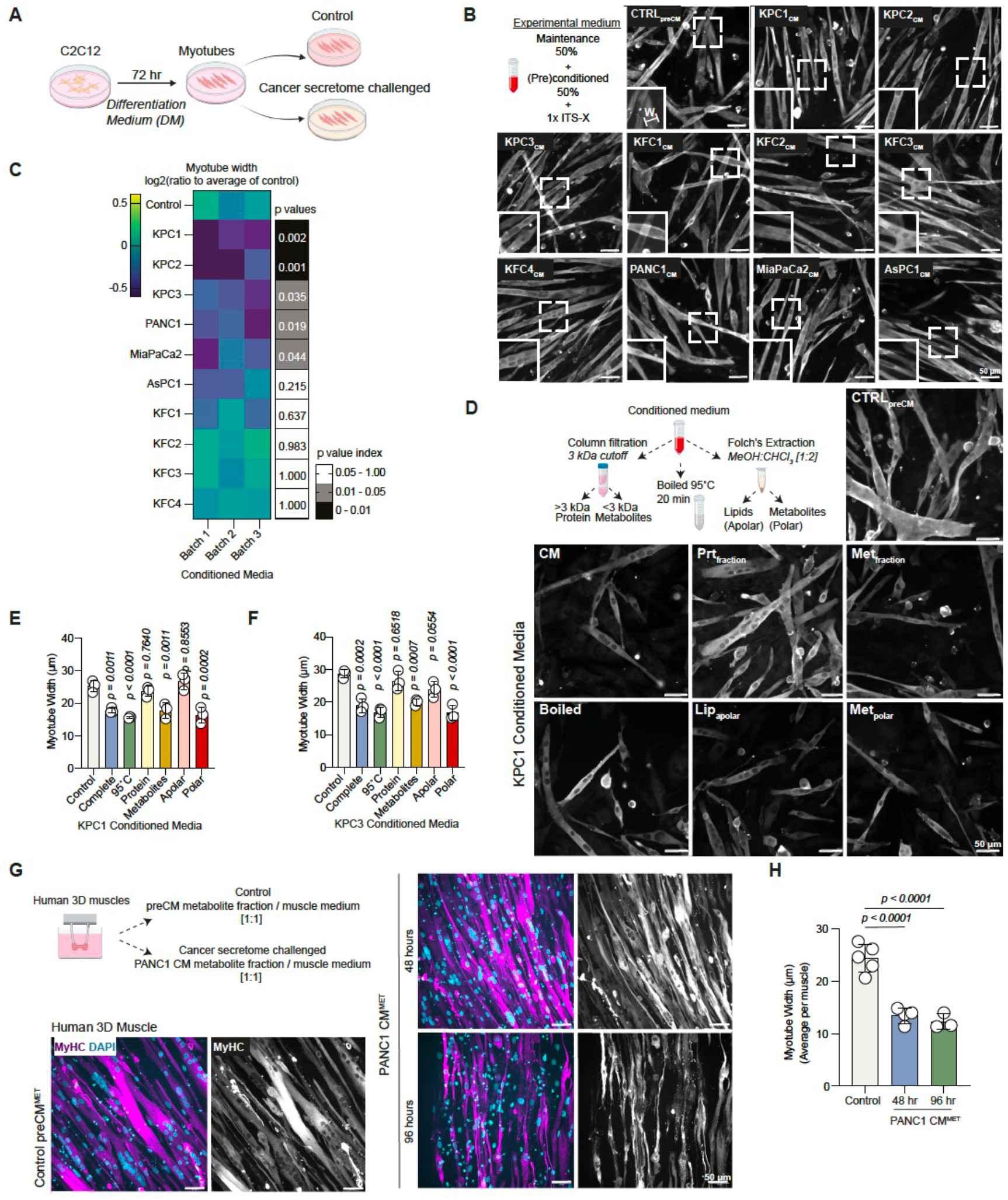
PDAC cell derived metabolites induce myotube atrophy. (A) Experimental scheme. (B) Scheme showing composition of experimental medium which was composed of 50% pre- or cancer cell conditioned medium diluted in myotube maintenance medium ensuring 1x of insulin-transferrin-selenium-ethanolamine (ITS-X) (top left) and representative images of MyHC-stained myotubes treated with conditioned media as indicated. Inlet; magnification of areas in dashed boxes. W: width, scale bars; 50 μm. (C) Heatmap, myotube width quantification from **(B)** as log2 ratio to average of controls. Grayscale heatmap indicates *P* values for each condition compared to control. Replicates are from 3 independently produced batches of conditioned media. (D) Experimental scheme and representative images of MyHC-stained myotubes treated with conditioned media fractions from KPC1 cells as indicated. Scale bars; 50 μm. (E-F) Width (μm) of myotubes treated with conditioned media fractions from KPC1 **(E)** and KPC3 **(F)** cells as indicated. Values are mean ± s.d. from average of >10 myotubes per replicate. (G) Experimental scheme and representative images of MyHC (magenta and grey right panel) and DAPI (blue)-stained 3D human engineered muscle tissues treated with control or PANC1-derived metabolites as indicated. Scale bars; 50 μm. (H) Width (μm) of myotubes within 3D engineered muscle tissues from **(G)**. Values are mean ± s.d. from average of >10 myotubes per replicate. C, E, F, H: Statistical significance was assessed with ordinary one-way analysis of variance (ANOVA) and Dunnett’s multiple comparisons test.

We identified myotubes with Myosin Heavy Chain (MyHC) staining and observed that myotube width was reduced 24 hours after exposure to conditioned media derived from KPC or human PDAC cells with p53 mutation, but not from PDAC cells with p53 loss or from C2C12-conditioned medium (Figure 1B & C). Myotubes exposed to KPC-conditioned medium also showed a modest reduction in the number of nuclei per myotube (Extended Data Fig. 1B), accompanied by an increase in the number of MyHC^+^ cells (Extended Data Fig. 1C), without a decrease in total MyHC^+^ area (Extended Data Fig. 1D), suggesting a mild defect in nuclei fusion. To characterise the PDAC-derived factor responsible for myotube atrophy, conditioned media were subjected to boiling to denature proteins, 3 kDa column filtration to separate protein (>3 kDa) and metabolite (<3 kDa) fractions, or Folch extraction to isolate lipids (apolar fraction) and polar metabolites (Figure 1D). Boiled conditioned media from KPC PDAC cells induced myotube atrophy at a similar rate to complete conditioned media, indicating that PDAC-secreted proteins are not responsible for this effect (Figure 1D-E) Indeed, the isolated protein fraction from KPC cells did not affect myotube morphology (Figure 1D-F). By contrast, the metabolite fraction of KPC cells induced atrophy to a similar extent as complete conditioned media, suggesting that the responsible factor is metabolite-derived (Figure 1D-F). Fractionation of metabolites indicated the polar fraction recapitulated the effects of complete and metabolite fraction of conditioned media, while the lipid fraction had no effect (Figure 1D-F). Interestingly, isolated polar metabolites from KFC-derived conditioned media were also sufficient to induce myotube atrophy, suggesting that protein components in the complete conditioned media may mask this metabolite-driven effect (Extended Data Fig. 1E & F).

Next, we sought to assess whether human myotubes are similarly sensitive to PDAC-derived metabolites. Immortalised human myoblasts were differentiated on ECM-coated surfaces *in vitro* and exposed to the metabolite fraction of conditioned media from human PANC1 cells. Following 24 hours of exposure, multinucleated myotube width was reduced (Extended Data Fig. 1G & H). To better recapitulate the complexity of skeletal muscle *in vitro*, we generated 3D engineered muscle tissues from the same human myoblasts^33^ and exposed them to PANC1-derived metabolites. Consistently, these metabolites induced atrophy in 3D human muscle constructs (Figure 1G & H), indicating that a metabolite-driven mechanism can induce atrophy in both human and mouse myofibres.

### PDAC derived metabolites increase myotube contractility

We next asked whether PDAC-derived metabolites actively induce atrophy or instead limits myotube growth and hypertrophy. Live imaging of myotubes from the onset of exposure to KPC1-derived metabolites showed that myotubes progressively decreased in width within the first hours of treatment (Figure 2A-C). RT-qPCR analysis of a panel of genes involved in proteolysis, autophagy, apoptosis, and myocyte differentiation and fusion showed an early induction of apoptosis-related genes, but no change in the expression of proteolysis-related genes 24 hours after exposure to KPC or KFC-derived metabolites (Extended Data Fig. 2A). Protein levels of FBXO32, a key mediator of atrophy, were only minimally increased at this timepoint (Extended Data Fig. 2B), suggesting that additional mechanisms contribute to the observed changes in myotube morphology. Live imaging suggested that nuclear movement within myotubes was less coordinated upon exposure to KPC-derived metabolites (Figure 2A), prompting us to analyse the migration of single nuclei within myotubes in detail. Nuclei in treated myotubes moved less directly (Figure 2D and Extended Data Fig. 2C & D), while their overall speed remained similar (Figure 2E and Extended Data Fig. 2E). These results indicate reduced coordination and alignment of nuclear movement across affected myotubes, suggesting cytoskeletal impairments may underlie myotube thinning. Staining for phosphorylated MLC2 at Ser19 revealed a strong increase in pMLC2 accumulation along actin stress fibres in the conditioned medium exposed myotubes (Figure 2F). Overall, the levels of pMLC2^Ser19^ were higher in myotubes exposed to KPC1-derived metabolites (Figure 2G). Since pMLC2 drives actin-myosin interactions and promotes stress fibre contraction^34^, these data suggest that myotube atrophy may be mechanically-mediated, arising from increased cytoskeletal tension and contractility.

**Figure 2:**
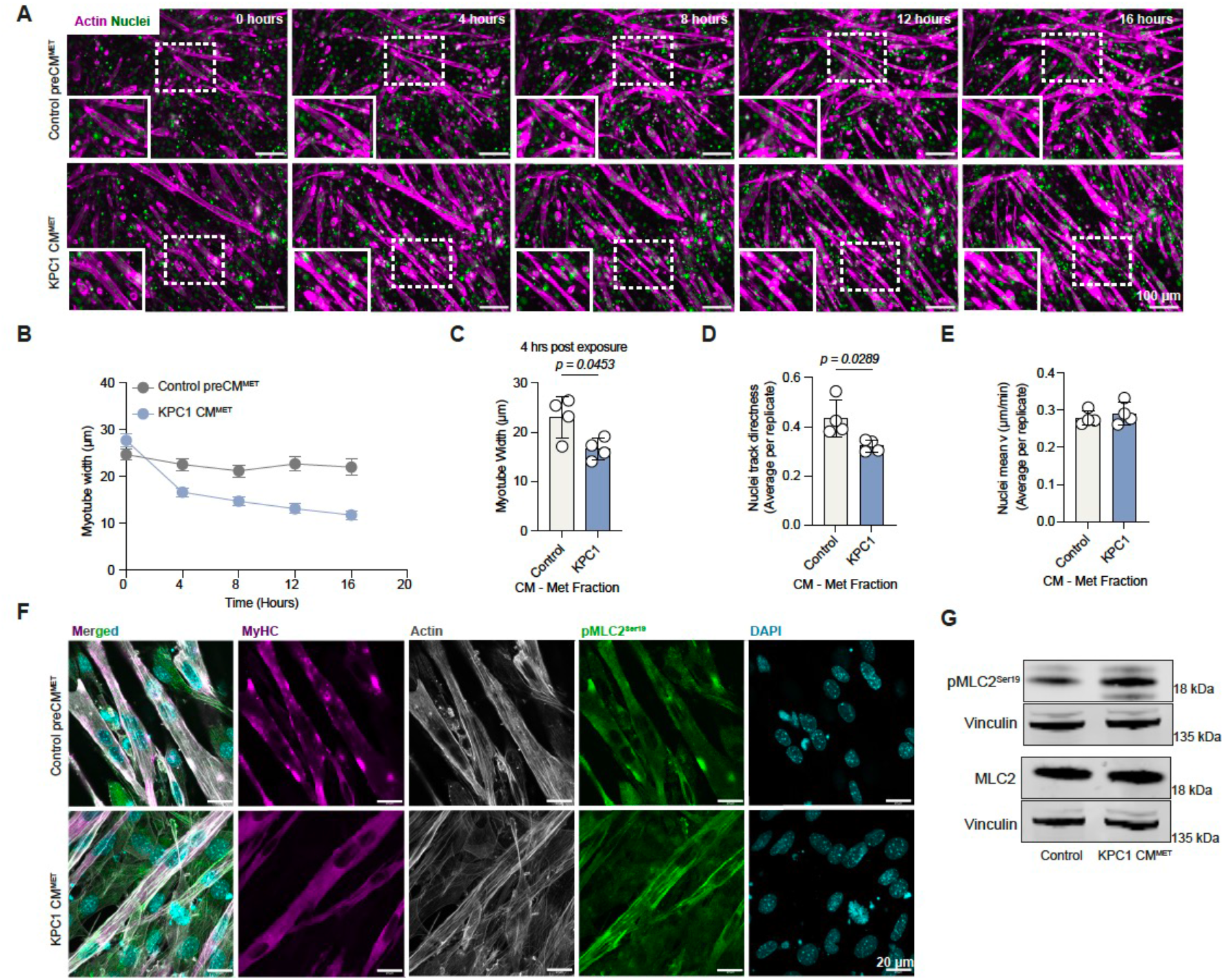
Myotube atrophy induced by PDAC cell-derived metabolites is defined by cytoskeletal stress through increased myosin-dependent contractility. (A) Representative images of Actin (magenta) and nuclei (green) labelled myotubes treated with Control or KPC1-derived metabolites over time as indicated. Inlet; magnification of areas in dashed boxes. Scale bars; 100 μm. (B) Myotube width (μm) over time from **(A)**. Values are mean+s.e.m. from four replicates per condition. (C) Myotube width (μm) from **(A)** 4 hours since beginning of treatment. Values are mean+s.d. from four replicates per condition. Statistical significance was assessed by two-tailed Welch’s *t*-test. (D-E) Myotube nuclei directness **(D)** and mean velocity (μm/min) **(E)** from **(A)**. Values are mean+s.d. from four replicates per condition. Statistical significance was assessed by two-tailed unpaired *t*-test. (F) Representative images of MyHC (magenta), Actin (grey), pMLC2^Ser19^ (green) and nuclei (cyan) stained myotubes treated with control or KPC1-derived metabolites. Scale bars, 20 μm. (G) Immunoblots of pMLC2^Ser19^, MLC2 and Vinculin (loading control) in C2C12 myotubes treated with Control or KPC1-derived metabolites. Blots are representative of three independent experiments. B-E: Values are average from >10 myotubes (B) or nuclei (C-E) per replicate and representative from three independent experiments.

### PDAC derived lactate induces myotube atrophy

To determine which metabolic feature derived from PDAC cells induces myotube atrophy, we performed gas chromatography-mass spectrometry (GC-MS) analysis of the metabolite fraction from conditioned media of two KPC and two KFC cell lines. Several metabolites were decreased compared to pre-conditioned media, including glucose, pyruvate and several amino acids. On the other hand, both KPC and KFC cells enriched their extracellular environment with high levels of lactate and most TCA cycle intermediates, including cis-aconitate, α-ketoglutarate, citrate, malate, succinate and fumarate (Figure 3A and S3A & B).

**Figure 3:**
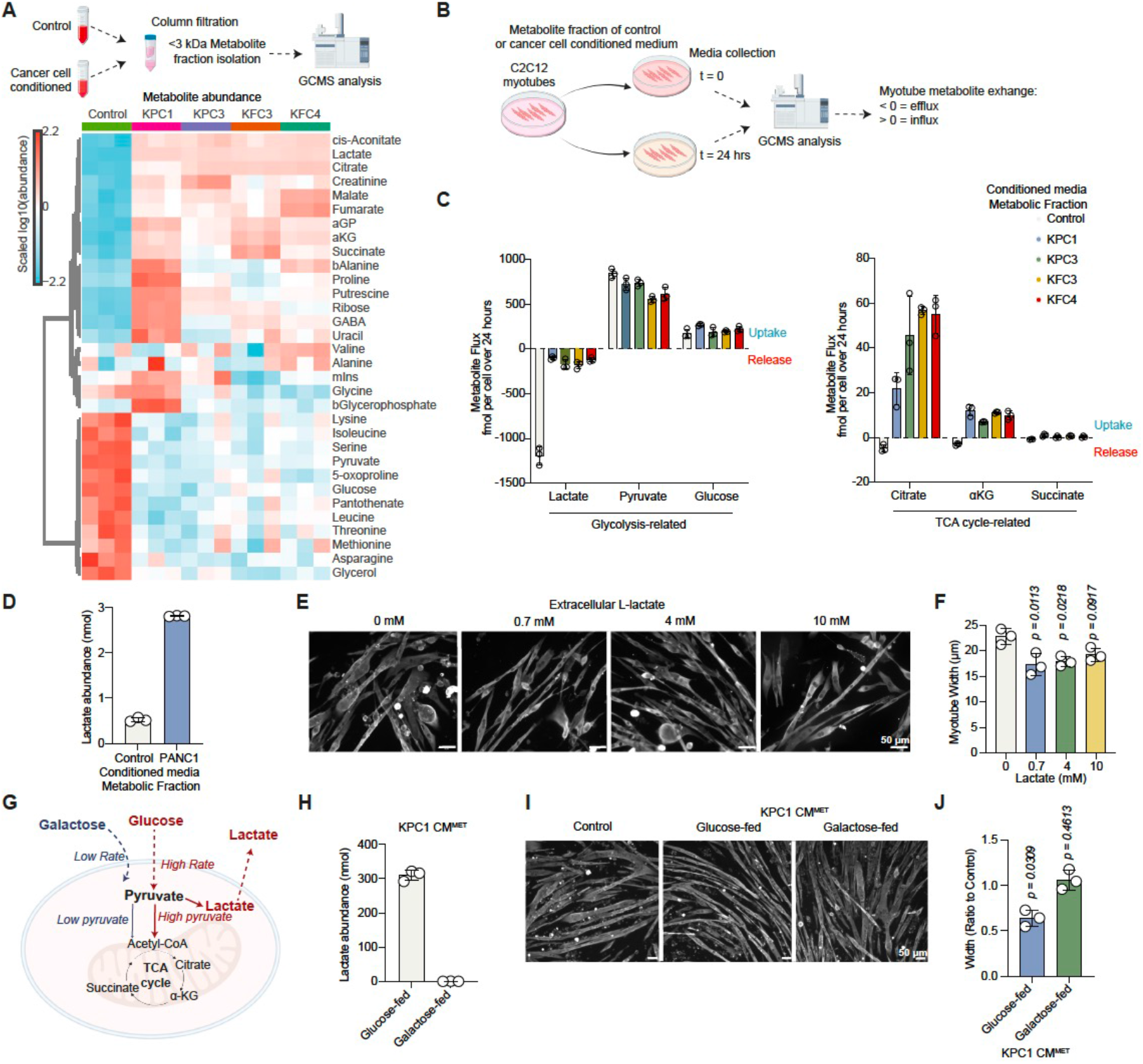
PDAC cell-secreted lactate induces myotube atrophy. (A) Top; Experimental scheme for quantification of metabolites in PDAC cell conditioned media. Bottom; Heatmap, abundance of detected metabolites in metabolite fraction from control or PDAC cell conditioned media. Replicates are from 3 independently produced conditioned media batches. (B) Schematic showing experimental strategy to quantify metabolite exchange in myotubes. (C) Metabolite flux (uptake or release, fmol per cell) of indicated glycolysis and TCA cycle related metabolites in myotubes treated with control or PDAC cell derived metabolites as indicated for 24 hr. Values are mean±s.d. from three replicates. (D) Lactate abundance (nmol) in metabolite fraction from control or PANC1 conditioned media. Values are mean±s.d. from three replicates. (E) Representative images of MyHC-stained myotubes treated with increasing concentrations of L-lactate as indicated. Scale bars; 50 μm. (F) Width (μm) of myotubes from **(E)**. Values are mean ± s.d. from n = 3 independent experiments with three replicates per experiment and >10 myotubes quantified per replicate. Statistical significance was assessed with ordinary one-way analysis of variance (ANOVA) and Tukey’s multiple comparisons test. (G) Schematic outlining utilisation of glucose or galactose as the main carbon donors. (H) Lactate abundance (nmol) in metabolite fraction of conditioned media from KPC1 cells supplemented with glucose or galactose. Values are mean±s.d. from three replicates. (I) Representative images of MyHC-stained myotubes treated with metabolite fraction from control, glucose or galactose-fed KPC1 cell conditioned media. Scale bars; 50 μm. (J) Width (μm) of myotubes from **(I)**. Values are mean ± s.d. and shown as ratio to control from n = 3 independent experiments with three replicates per experiment and >10 myotubes quantified per replicate. Statistical significance was assessed with two-tailed one-sample *t*-test on natural log transformed values.

We then analysed media from myotubes exposed to control or cancer cell derived metabolites acquired at the onset of exposure (t = 0) and after 24 hours (t = 24 hours) to assess changes in metabolite exchange rates (Figure 3B and S3C). We initially observed that - compared to control medium treated cells - conditioned medium exposed myotubes largely stopped lactate efflux, while glucose or pyruvate influx remained unchanged (Figure 3C). These myotubes also appeared to uptake TCA intermediates from the media, most prominently citrate and αKG, although their overall abundance was lower compared to other metabolites (Figure 3C and S3C). Additionally, uptake of serine and export of glycine were modestly reduced (Extended Data Fig. 3D). Collectively these data identify lactate as the most impacted metabolite. While lactate is normally exported by myotubes in control medium, but exposure to the lactate-rich PDAC media blocks this efflux. Consistently, PANC-1-derived conditioned media is also enriched in lactate (Figure 3D).

To examine the impact of lactate directly, we exposed myotubes to increasing concentrations of extracellular lactate and assessed their morphology. Lactate administration at 0.7 mM led to thinner myotubes, and while higher doses did not produce an additive effect (Figure 3E & F), increasing lactate levels also did not proportionally raise intracellular lactate levels (Extended Data Fig. 3E & F). It is worth noting that 0.7 mM L-lactate matches the concentration of lactate detected in conditioned media (Extended Data Fig. 3G). To test whether lactate is the main driver of myotube atrophy in PDAC-conditioned media, we generated media from PDAC cells with galactose or glucose as the carbon source. Galactose metabolism is slower than glucose, generating less pyruvate which is directed almost exclusively to the mitochondria, resulting in reduced pyruvate conversion to lactate and lower lactate secretion^35^ (Figure 3G). Consistently, conditioned media from galactose-fed KPC1 cells did not contain lactate. Proline and alanine levels were also decreased in the conditioned medium from galactose-fed compared to glucose-fed cells, while other metabolites remained unchanged (Figure 3H and Extended Data Fig. 3H-J). The metabolite fraction from galactose-derived conditioned media did not induce myotube atrophy (Figure 3I &J), supporting the model that elevated lactate is a primary metabolic driver of PDAC-induced atrophy. Interestingly, whereas lactate was initially considered a ‘metabolic waste’ of increased glycolysis during exercise contributing to muscle fatigue, it is emerging as an energy substrate and signalling molecule with pleiotropic roles in muscle physiology^36^. In MCT1 expressing muscle, lactate can fuel the TCA cycle increasing the abundance of oxidative over glycolytic fibres^37^. Increasing muscle or circulating lactate has also been associated with cancer cachexia^38–41^. Therefore, we aimed to assess intracellular lactate levels and understand its implications.

### PDAC affected myotubes exhibit reduced NAD^+^/NADH ratio and lower αKG synthesis

Lactate flux across the plasma membrane is driven by its concentration as well as pH gradients. Typically, under normal conditions, increased lactate production promotes lactate secretion when extracellular lactate levels are low. However, upon equimolar concentrations or at higher extracellular lactate concentrations, lactate flux stalls or gets reversed respectively^42^ (Figure 4A). Myotubes exposed to PDAC-cell derived metabolites for 24 hours had elevated intracellular lactate levels (Figure 4B), while glucose and pyruvate were unchanged (Extended Data Fig. 4A & B). In addition, the intracellular lactate to pyruvate ratio was increased in conditioned medium exposed myotubes (Figure 4C). To investigate whether intracellular lactate accumulation was a result of increased synthesis we administered [U-^13^C_6_]-D-glucose 20 hours after treatment onset for 4 hours and traced glucose-derived carbon incorporation into downstream metabolites. Myotubes exposed to PDAC cell derived metabolites showed reduced glucose-derived pyruvate and lactate synthesis (Figure 4D and Extended Data Fig. 4C), suggesting that intracellular lactate accumulation arises from slower lactate usage and impaired export.

**Figure 4:**
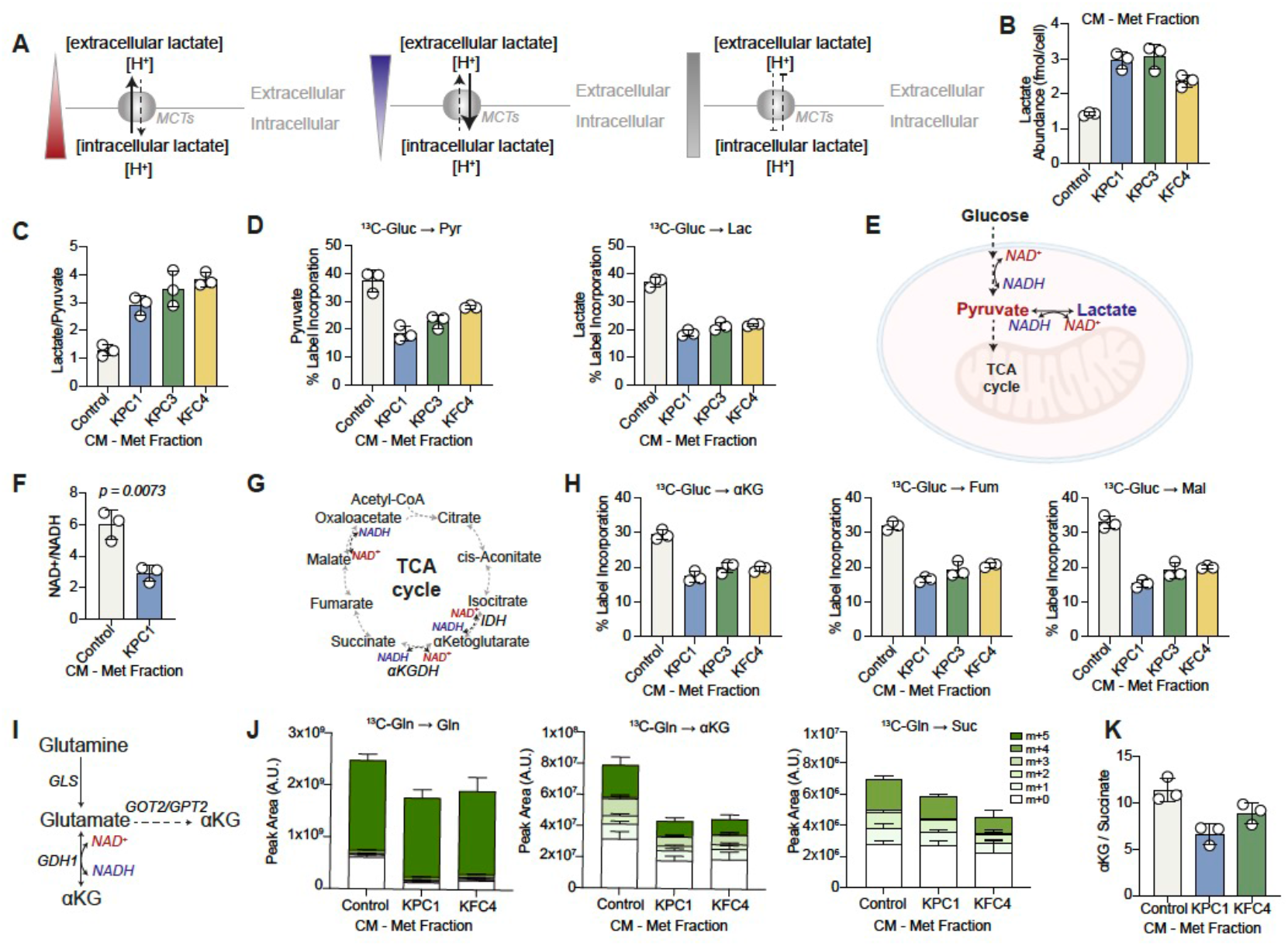
PDAC cell derived metabolites impact NAD^+^/NADH balance and decrease αKG levels in myotubes. (A) Schematic depicting lactate transport and its dependence on concentration gradients. (B) Lactate abundance (fmol/cell) in myotubes treated with control or PDAC cell derived metabolites as indicated. Values are mean±s.d. from three replicates. (C) Intracellular lactate to pyruvate ratio in myotubes treated with control or PDAC cell derived metabolites as indicated. Values are mean±s.d. from three replicates. (D) Incorporation (total label %) of [U-^13^C_6_]-D-glucose-derived, labelled carbons into pyruvate (left) and lactate (right) in myotubes treated with control or PDAC cell derived metabolites as indicated. Values are mean±s.d. from three replicates. (E) Simplified schematic showing NAD^+^/NADH balance correlation with lactate/pyruvate abundance. (F) NAD^+^/NADH ratio in myotubes treated with control or PDAC cell derived metabolites as indicated. Values are mean±s.d. and average of triplicate wells from three independent experiments. Statistical significance was assessed with two-tailed unpaired *t*-test. (G) Schematic of TCA cycle dependence on NAD^+^/NADH balance. (H) Incorporation (total label %) of [U-^13^C_6_]-D-glucose-derived, labelled carbons into αKG (left), fumarate (middle) and malate (right) in myotubes treated with control or PDAC cell derived metabolites as indicated. Values are mean±s.d. from three replicates. (I) Simplified schematic of glutamine to αKG reaction and dependence on NAD^+^/NADH balance. (J) Incorporation of [U-^13^C_5_]-L-glutamine-derived, labelled carbons into glutamine (left), αKG (middle) and succinate (right) in myotubes treated with control or PDAC cell derived metabolites as indicated. Values are mean±s.d. from three replicates. (K) αKG over succinate ratio in myotubes treated with control or PDAC cell derived metabolites as indicated. Values are mean±s.d. from three replicates.

The cytosolic redox state can be inferred from the relative abundance of lactate over pyruvate, which reflects the NAD^+^/NADH ratio assuming near equilibrium of LDH activity^43,44^ (Figure 4E). Consistently, the NAD^+^/NADH ratio was reduced in myotubes exposed to KPC1-derived metabolites (Figure 4F).

NAD^+^ is required for key oxidative steps of the TCA cycle, including reactions catalysed by isocitrate dehydrogenase (IDH) and α-ketoglutarate dehydrogenase (αKGDH), suggesting that decreased NAD^+^ availability may constrain oxidative TCA flux (Figure 4G). Tracing [U-^13^C_6_]-D-glucose-derived carbon incorporation into TCA intermediates supported decreased glucose to the cycle (Figure 4H). While intracellular α-ketoglutarate (αKG) levels were reduced (Extended Data Fig. 4D), fumarate (Extended Data Fig. 4E) and malate (Extended Data Fig. 4F) remained unchanged, indicating a specific perturbation at the level of αKG. aKG can also be generated from glutamine *via* glutamate, through either an NAD^+^ dependent oxidative reaction or transamination reactions^45^ (Figure 4I). To test whether myotubes switch towards glutamine dependency to fuel oxidative phosphorylation, we administered [U-^13^C_5_]-L-glutamine and traced its incorporation into TCA metabolites. Although glutamine uptake was similar, downstream labelling was reduced, most prominently for αKG, and to a lesser extent, succinate, while citrate and other intermediates showed unchanged labelling (Figure 4J and Extended Data Fig. 4G). Consistently, the αKG/succinate ratio was decreased, suggesting reduced αKG availability relative to downstream intermediates.

Glutamine can also support the TCA cycle *via* reductive carboxylation, where αKG is converted to citrate through reversal of oxidative TCA flux ^46^. The citrate m+5/m+4 ratio was unchanged between conditions, suggesting no increase in reductive carboxylation flux (Extended Data Fig. 4H). Given the reduced αKG pool and the preserved citrate labelling, the relative contribution of reductive versus oxidative glutamine metabolism appears maintained; despite decreased entry at the αKG step, downstream flux to citrate appears preserved. Levels of 2-hydroxyglutarate (2HG), which is derived from αKG, were also decreased (Extended Data Fig. 4I), consistent with lower αKG pools, while the ratio of labelled 2HG to αKG remained unchanged (Extended Data Fig. 4J), indicating that there is no alteration in 2HG generation rates between conditions. Serine metabolism, which can also contribute to cellular redox balance^47^, was also perturbed, with reduced serine levels (Extended Data Fig. 4K) and decreased glucose-derived labelling into glycine (Extended Data Fig. L & M). Overall, these findings suggest that decreased NAD^+^ availability constraints oxidative metabolism and selectively limits αKG production, positioning αKG as a key metabolic node affected in myotubes exposed to PDAC-derived metabolites.

### Restoration of NAD^+^ levels rescues myotube atrophy

Next, we asked whether exposure to PDAC-derived metabolites alters cellular energy balance, redox homeostasis and mitochondrial bioenergetics in myotubes. Exposure to PDAC-derived metabolites increased AMP accumulation and AMP/ATP ratio suggesting a modest energetic stress, but without affecting total ATP levels or the ADP/ATP ratio (Extended Data Fig. 5A-D). Affected myotubes showed decreased intracellular GSH levels (Extended Data Fig. 5E) and lower GSH/GSSG ratio (Figure 5A), indicating increased oxidative stress. Consistently, cytosolic ROS levels were increased (Figure 5B), whereas mitochondrial superoxide levels were unchanged (Extended Data Fig. 5F). Myotubes had similar mitochondrial content and membrane potential between conditions (Extended Data Fig. G & H).

**Figure 5:**
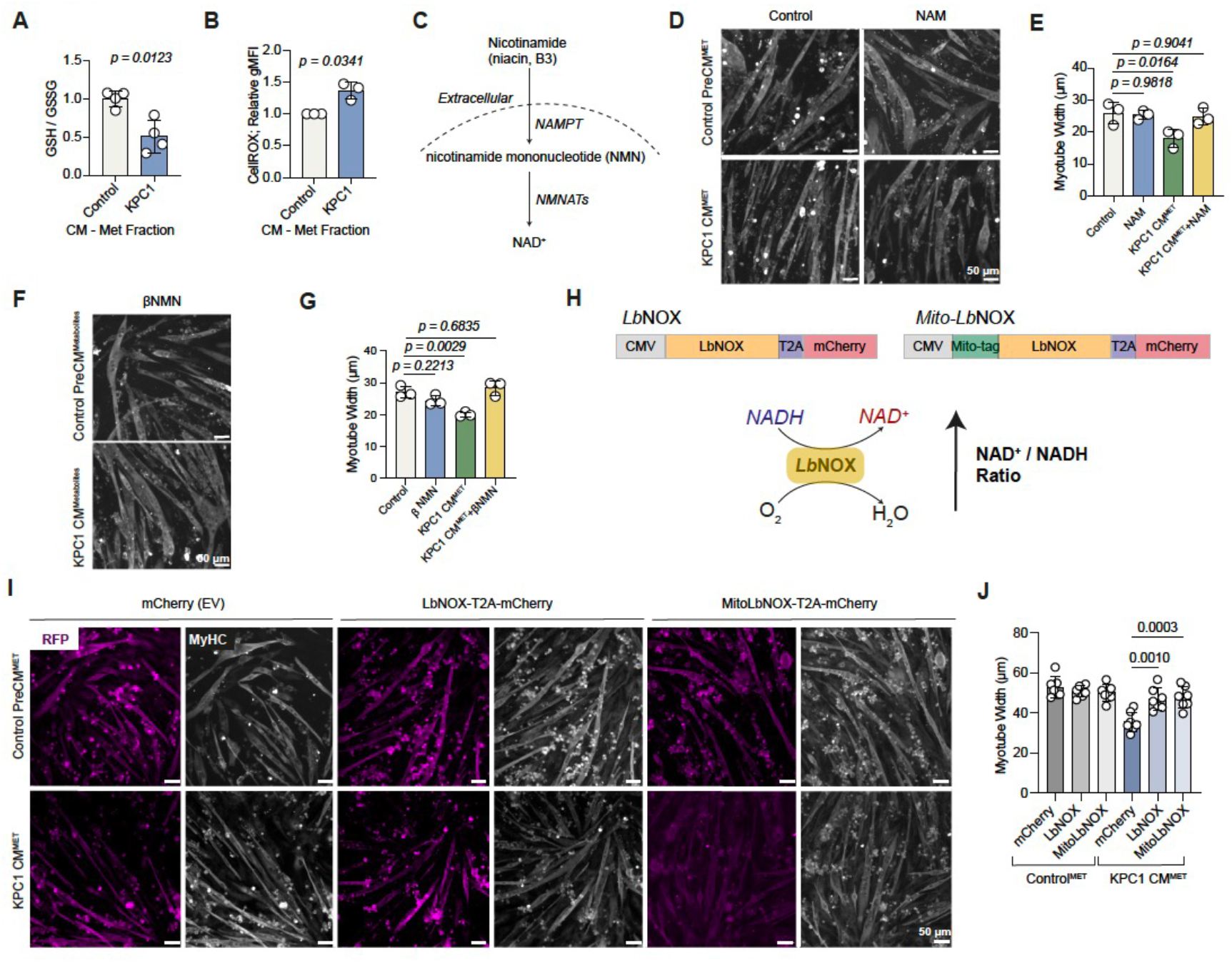
NAD^+^/NADH imbalance defines PDAC-induced myotube atrophy. (A) GSH/GSSG ratio in myotubes treated with control or KPC1 cell derived metabolites. Values are mean±s.d. and average of >3 wells from four independent experiments. Statistical significance was assessed with two-tailed Welch’s *t*-test. (B) Cellular ROS (CellROX, gMFI) levels of myotubes treated with control or KPC1 cell derived metabolites as indicated. Values are mean±s.d. and average of triplicate wells from three independent experiments. Statistical significance was assessed with two-tailed one-sample *t*-test on natural log transformed values. (C) Simplified schematic depicting nicotinamide catabolism. (D) Representative images of MyHC-stained myotubes treated with metabolite fraction from control or KPC1 cell conditioned media ± nicotinamide (NAM). Scale bars; 50 μm. (E) Width (μm) of myotubes from **(D)**. Values are mean ± s.d. and average of three replicates with >10 myotubes quantified per replicate from three independent experiments. Statistical significance was assessed with ordinary one-way analysis of variance (ANOVA) and Dunnett’s multiple comparisons test. (F) Representative images of MyHC-stained myotubes treated with metabolite fraction from control or KPC1 cell conditioned media supplemented with nicotinamide mononucleotide (βNMN). Scale bars; 50 μm.(G) Width (μm) of myotubes from **(F)**. Values are mean ± s.d. from three replicates with >10 myotubes quantified per replicate. Statistical significance was assessed with ordinary one-way analysis of variance (ANOVA) and Dunnett’s multiple comparisons test. (H) Simplified schematic showing NAD+ regeneration from *Lb*NOX and construct design. (I) Representative images of mCherry (magenta) and MyHC (grey)-stained myotubes expressing mCherry (empty vector, EV), LbNOX-T2A-mCherry or MitoLbNOX-T2A-mCherry and treated with metabolite fraction from control or KPC1 cell conditioned media. Scale bars; 50 μm. (J) Width (μm) of myotubes from **(I)**. Values are mean ± s.d. from two independent experiments with >3 replicates per experiment and >10 myotubes quantified per replicate. Statistical significance was assessed with ordinary one-way analysis of variance (ANOVA) and Tukey’s multiple comparisons test.

To determine whether redox alterations contribute to atrophy, we exposed control and PDAC-metabolite exposed myotubes to a panel of antioxidants, including NAC, cell permeable glutathione (GSH-EE) and Trolox, the mitochondrial superoxide scavenger mitoTEMPO, the ferroptosis inhibitors ferrostatin-1 and liproxstatin-1, as well as nicotinamide to restore NAD^+^ levels. Among these, only nicotinamide rescued PDAC metabolite-induced atrophy (Figure 5C-E and Extended Data Fig. 5I & J), suggesting that restoring NAD^+^ availability, rather than antioxidant treatment is sufficient to reverse this phenotype. Consistently, administration of the NAD^+^ precursor, NMN also rescued myotube atrophy (Figure 5F & G).

Affected myotubes showed a modest but consistent increased expression of complex I subunits, reflecting a compensatory response to impaired NAD^+^ availability and mitochondrial oxidative phosphorylation capacity (Extended Data Fig. 5K & L). Myotubes expressing *Lactobacillus brevis* NADH oxidase (*LbNOX*) or its mitochondrial-targeted variant (mito-*LbNOX*), which regenerate NAD^+^ through NADH oxidation using molecular oxygen (Figure 5H), were resistant to PDAC-derived metabolite induced atrophy^48^ (Figure 5I,J). Overall, these findings underpin that NAD^+^ homeostasis is a key determinant of myotube morphology, although associated alterations in redox balance are not sufficient to drive atrophy.

### αKG directs myotube morphology

To determine whether reduced αKG levels, resulting from NAD^+^ limitation in affected myotubes, contribute to atrophy we considered its role beyond a TCA cycle substrate. αKG acts as a co-substrate for a large family of dioxygenase enzymes, including the Jumonji-domain histone demethylases. These regulate chromatin by converting αKG to succinate, with the latter also being a competitive inhibitor of the reaction^49^. Myotube differentiation and muscle remodelling rely on dynamic alterations of trimethylation of histone 3 lysine 27 (H3K27me3). H3K27me3 is a repressive mark and is typically removed from chromatin regions to allow transcription of differentiation and lineage genes such as *Myog*, while being enriched to repress genes associated with proliferation^50,51^. H3K27me3 demethylation is mediated by JMJD3, an αKG dependent enzyme (Figure 6A).

**Figure 6:**
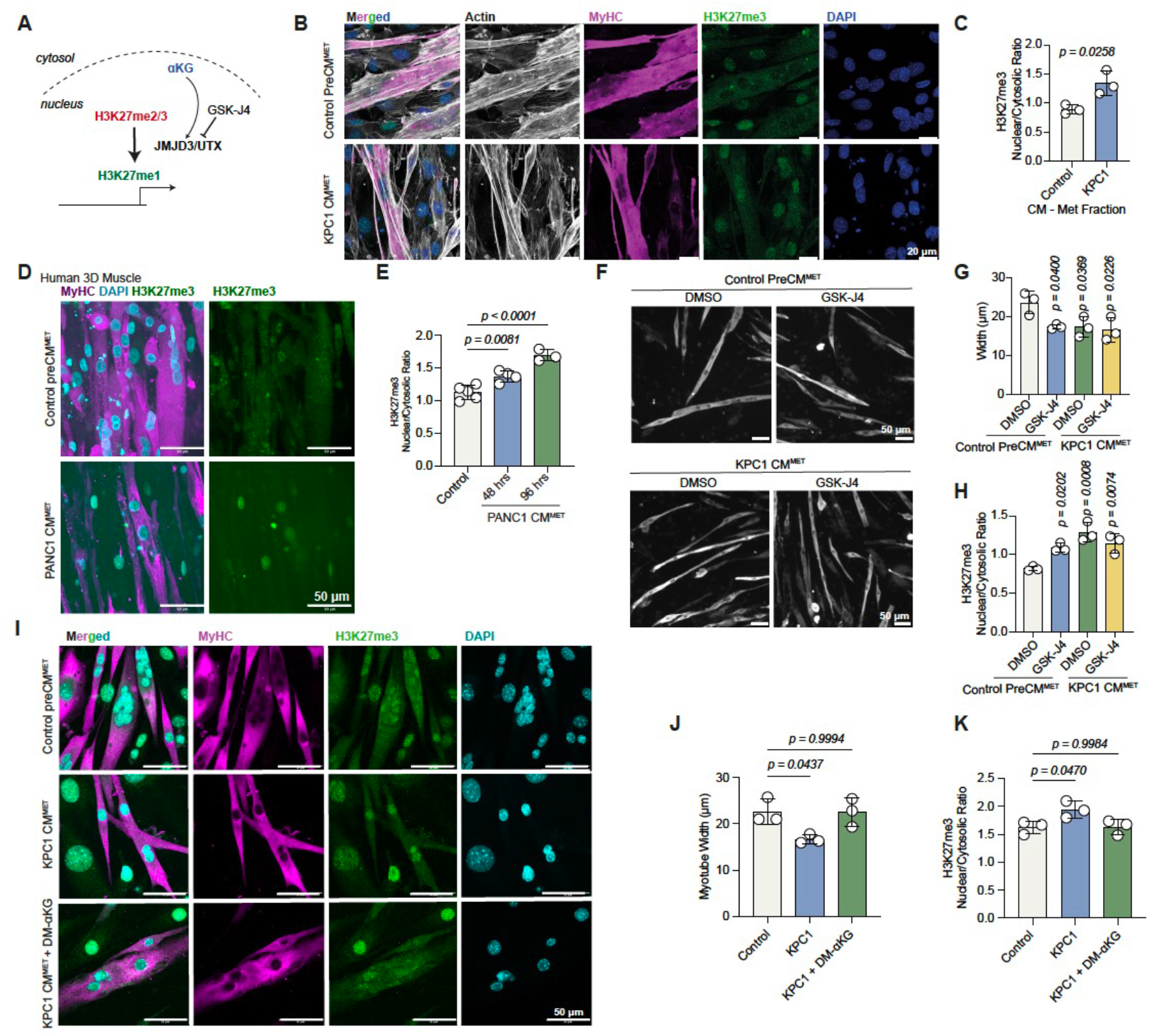
αKG levels in PDAC-induced myotube atrophy influence histone methylation. (A) Simplified schematic outlining αKG requirement for histone demethylation. (B) Representative images of Actin (grey), MyHC (magenta), H3K27me3 (green) and DAPI-stained myotubes treated with metabolite fraction from control or KPC1 cell conditioned media. Scale bars; 20 μm. (C) H3K27me3 nuclear to cytosolic ratio of myotubes from **(B)**. Values are mean±s.d. and average of >3 replicates from three independent experiments. Statistical significance was assessed with two-tailed unpaired *t*-test. (D) Representative images of MyHC (magenta), H3K27me3 (green) and DAPI (blue)-stained 3D human muscle constructs treated with control or PANC1-derived metabolites as indicated. Scale bars; 50 μm. (E) H3K27me3 nuclear to cytosolic ratio of myotubes from **(D)**. Values are mean ± s.d. from average of >10 myotubes per replicate. (F) Representative images of MyHC-stained myotubes treated with metabolite fraction from control or KPC1 cell conditioned media and either DMSO or GSK-J4. Scale bars; 50 μm. (G) Width (μm) of myotubes from **(F)**. Values are mean ± s.d. and average of >3 replicates from three independent experiments. (H) H3K27me3 nuclear to cytosolic ratio of myotubes from **(F)**. Values are mean ± s.d. and average of >3 replicates from three independent experiments. (I) Representative images of MyHC (magenta), H3K27me3 (green) and DAPI (blue)-stained myotubes treated with control and KPC1-derived metabolites ±DM-αKG as indicated. Scale bars; 50 μm. (J) Width (μm) of myotubes from **(I)**. Values are mean ± s.d. from three replicates. (K) H3K27me3 nuclear to cytosolic ratio of myotubes from **(I)**. Values are mean ± s.d. from three replicates. E, G, H, J, K: Statistical significance was assessed with ordinary one-way analysis of variance (ANOVA) and Dunnett’s multiple comparisons test.

Myotubes exposed to PDAC-metabolites exhibited elevated levels of nuclear H3K27me3 (Figure 6B & C). This was consistent in human myotubes exposed to PANC1-derived metabolites either in 2D (Extended Data Fig. 6A & B) or 3D muscle constructs (Figure 6D & E). Lactate-treated myotubes displayed reduced αKG levels and decreased αKG to fumarate ratio (Extended Data Fig. 6C & D). This was accompanied by increased nuclear H3K27me3 (Extended Data Fig. 6E). To test whether αKG-dependent H3K27me3 demethylation contributes to atrophy, we used the JMJD3 inhibitor GSK-J4, which increased H3K27me3 and reduced myotube width, but did not further enhance atrophy in PDAC-exposed myotubes (Figure 6F-H). Conversely, administration of cell-permeable αKG rescued myotube width and restored H3K27me3 to control levels (Figure 6I-K) in PDAC-metabolite exposed myotubes.

Collectively these findings suggest that reduced αKG is a signature of atrophic myotubes and may contribute in atrophy, at least in part, by limiting αKG-dependent chromatin remodelling.

### Muscle of cachectic mice has reduced αKG

To assess whether these metabolic signatures are recapitulated *in vivo*, we turned to mouse models. Modelling cancer cachexia in mice presents several challenges. In contrast to human disease, muscle loss is often modest and typically occurs only at very advanced stages ^5,52^. This discrepancy may also be influenced by the relatively young age of experimental mice, which are still in an anabolic state and may therefore partially compensate for muscle loss through ongoing growth or regeneration^53,54^. Moreover, in some models, rapid tumour progression leads to humane endpoints before significant body weight loss is observed^55^.

To address these limitations, we employed two complementary mouse models. First, we used immunocompetent C57BL/6 mice subcutaneously implanted with KPC1 cells. After two weeks, when mice were collected due to tumour burden, they displayed features of early cachexia, with preserved body weight but modest reduction primarily in adipose tissue and to a lesser degree in muscle (Figure 7A-D and Extended Data Fig. 7A-C). Second, we utilised a model of severe cachexia in which intranasal administration of Ad5CMV-Cre induces non–small cell lung cancer driven by Kras-G12D and Lkb1 loss (KL model). This model progresses over a longer time course, allowing the development of overt cachexia, including significant body weight loss, as well as pronounced adipose and muscle wasting at endpoint (Figure 7A, E-G and Extended Data Fig. 7D-F). In KL mice, gastrocnemius myofibres were reduced in size, without switching to slow myosin fibre type and showed modest increase in *Fas*, *Fbxo32*, and *Fbxo31* expression (Figure 7H & I and Extended Data Fig. 7G & H).

**Figure 7:**
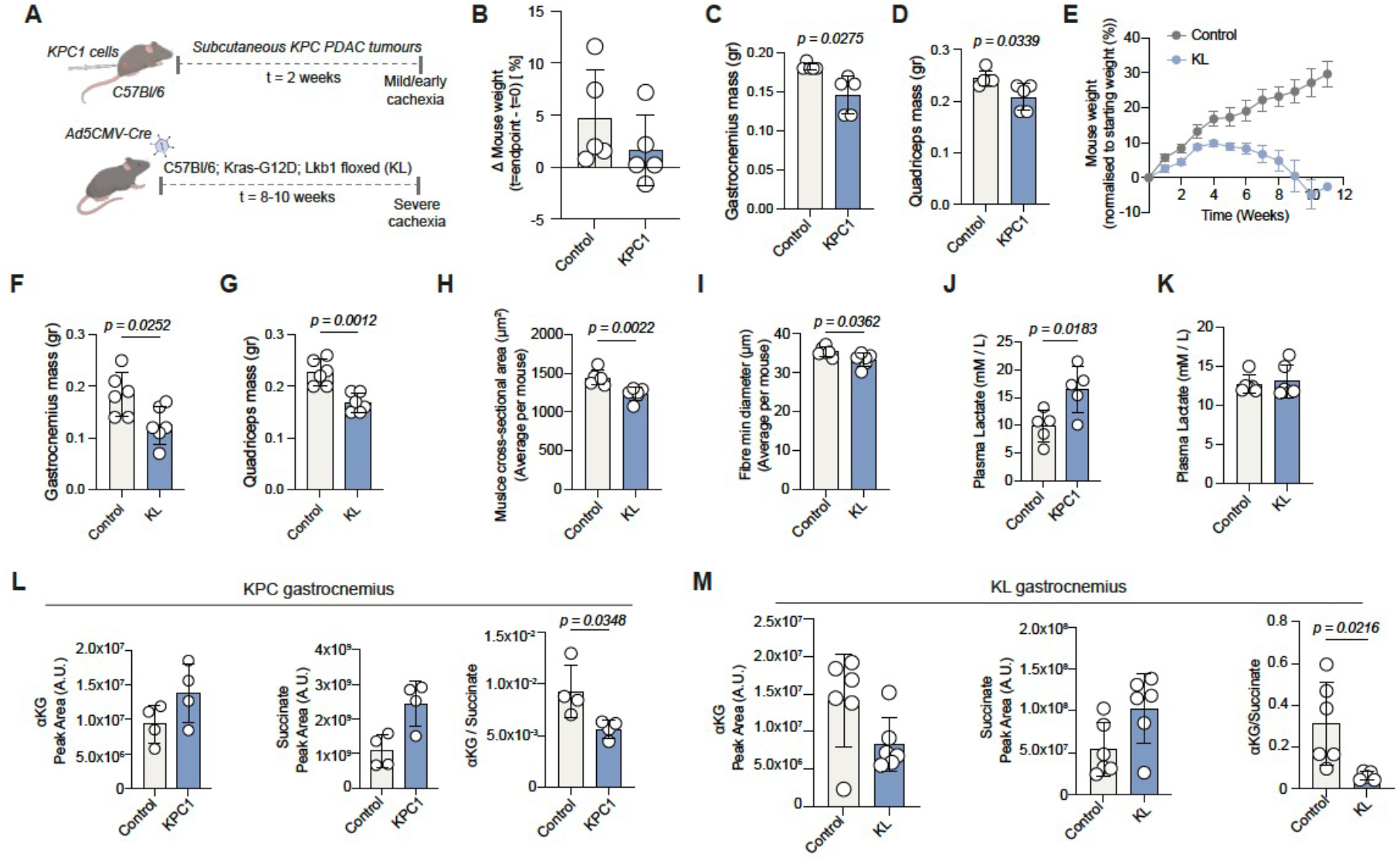
Muscle exhibits reduced αKG/succinate ratio in cancer-associated cachexia *in vivo.* (A) Schematic of mouse models used to model ‘early’ and ‘severe’ cachexia. (B) Weight difference (%) of endpoint (minus tumour weight) vs starting weight (prior tumour implantation) in KPC1 model. Values are mean±s.d. from n = 5 control and n = 5 KPC1 transplanted mice. (C-D) Gastrocnemius **(C)** and quadricep **(D)** muscle mass of mice from **(B)**. Values are mean±s.d. from n = 5 control and n = 5 KPC1 transplanted mice. (E) Weight trajectories of control or KL mice since Cre induction. Values are mean±s.e.m. from n = 6 control and n = 6 KL mice. (F-G) Gastrocnemius **(F)** and quadricep **(G)** muscle mass of mice from **(E)**. Values are mean±s.d. from n = 6 control and n = 6 KL mice. (H-I) Gastrocnemius muscle fibre cross-sectional area (μm^2^) **(H)** and min diameter (μm) **(I)** of mice from **(E)**. Values are mean±s.d. from n = 6 control and n = 6 KL mice. (J) Plasma lactate levels (mM/L) in control or KPC1 mice. Values are mean±s.d. from n = 7 control and n = 7 KPC1 transplanted mice. (K) Plasma lactate levels (mM/L) in control or KL mice. Values are mean±s.d. from n = 5 control and n = 5 KL mice. (L) Abundance of αKG (left), succinate (middle) and αKG to succinate ratio in gastrocnemius muscle of control or KPC1 mice. Values are mean±s.d. from n = 4 control and n = 4 KPC1 transplanted mice. (M) Abundance of αKG (left), succinate (middle) and αKG to succinate ratio in gastrocnemius muscle of control or KL mice. Values are mean±s.d. from n = 6 control and n = 6 KL mice. C, D, F, G, H, I: Statistical significance was assessed with two-tailed Welch’s *t*-test. J, L, M: Statistical significance was assessed with two-tailed unpaired *t*-test.

KPC1 tumour-bearing mice exhibited elevated circulating lactate levels (Figure 7J). It has been previously reported that cachexia resulting from lung tumours in mice and humans is a result of elevated lactate levels in the circulation^56^. Lkb1 loss in lung adenocarcinoma correlates with increased circulating lactate levels^38^, but in this study KL mice didn’t present increased circulating lactate at endpoint (Figure 7K).

To assess muscle metabolic alterations, we performed liquid chromatography–mass spectrometry (LC– MS)-based metabolomic profiling of gastrocnemius muscle from KPC1 and KL mice (Extended Data Fig. 7I). The gastrocnemius of KPC1 mice presented with decreased αKG/succinate ratio (Figure 7L), while circulating αKG levels also were moderately decreased (Extended Data Fig. 7J & K). In KL mice with advanced cachexia and muscle loss, αKG levels were reduced in both the circulation and gastrocnemius muscle, accompanied by a diminished αKG/succinate ratio (Figure 7M and Extended Data Fig. 7L & M). Collectively, these findings suggest that disrupted αKG homeostasis is a conserved feature of cancer-associated muscle wasting *in vivo* and may contribute to atrophy.

## DISCUSSION

Cancer cachexia has long been regarded as a secondary, paraneoplastic condition. However, it is increasingly recognised as complex and multifactorial disease with profound consequences for both patient survival and tumour progression. Despite its clinical impact, there are currently no effective therapeutic strategies^8,57^. This is in part due to the heterogeneous and systemic nature of the syndrome, as well as the lack of experimental models that fully recapitulate its origins and progression limiting our mechanistic understanding. In this study, we aimed to address these challenges by combining reductionist *in vitro* systems with complementary *in vivo* models to uncover mechanistic insights into tumour-muscle crosstalk.

Our findings define a metabolite-driven mechanism of cancer-associated muscle wasting, in which PDAC-derived lactate disrupts metabolic homeostasis in skeletal muscle. Previous work also noted an ability of lactate to decrease myotube width^32^. Elevated extracellular lactate alters lactate flux, leading to intracellular accumulation and a reduction in the NAD⁺/NADH ratio. Consistent with previous reports linking NAD⁺ depletion to mitochondrial dysfunction and cachexia severity^58–60^, we show that this redox imbalance limits αKG availability, thereby impairing αKG-dependent reactions, including chromatin remodelling. Previous studies have also demonstrated a role for NAD+ dependent regulation of the epigenetic regulator SIRT1 in muscle development^61^. In parallel, we identify increased pMLC2^Ser19^ as a downstream consequence of PDAC-associated cachexia, suggesting that metabolic stress is coupled to cytoskeletal tension to drive myotube atrophy. In smooth muscle, contractility and shortening can be mediated by increased MLC2 phosphorylation under RhoA and ROCK signalling^62,63^.

These findings are consistent with emerging evidence that lactate is not merely a metabolic by-product but an active signalling molecule in cachexia. Recent work has shown that tumour-derived lactate is sufficient to drive adipose and muscle wasting *in vivo*. Lactate was shown to induce lipolysis *via* GPR81-dependent pathways^56^. The activation of the G-protein-coupled receptor GPR81 by lactate activated a RhoA–ROCK1–p38 pathway in adipocytes, which drives adipose tissue browning and lipolysis contributing to weight loss^56^. It has also been recently discovered that lactate can also directly regulate gene expression through histone lactylation^64^. On the other hand, pyruvate administration has been shown to prevent the onset of cachectic features in myotubes *via* STAT3 signalling^65^. Our data suggests that the relative abundance of lactate over pyruvate has important metabolic implications in muscle cells and extend these observations by demonstrating that lactate can also act through redox-mediated depletion of NAD⁺ and αKG; metabolic alterations that may act through chromatin regulation and cytoskeletal dynamics in muscle cells.

We show that restoration of NAD⁺ or αKG levels rescues myotube atrophy, highlighting a potential therapeutic axis. This is supported by studies demonstrating that NAD⁺ replenishment through niacin supplementation improves mitochondrial function and attenuates cachexia phenotypes in preclinical models^58^, and that αKG supplementation can preserve muscle function and reduce frailty *in vivo*^66,67^. Systemic NAD^+^ deficiency has been reported in adult-onset mitochondrial myopathy patients. Niacin restored NAD^+^ levels and rescued muscle strength and mitochondrial biogenesis^68^. αKG which can be generated during endurance exercise, attenuated corticosterone-induced protein degradation and rescued muscle atrophy in a Duchenne muscular dystrophy mouse model. In this model, αKG Inhibited the expression of proline hydroxylase 3 (PHD3) and its interaction with the β2 adrenergic receptor (ADRB2), increasing ADRB2 stability^67^. Together, these findings position the NAD⁺-αKG axis as a tractable metabolic vulnerability in cancer-associated muscle wasting.

We argue that the integration of *in vitro* systems with *in vivo* models is essential to resolve mechanistic pathways that may be masked in whole-organism settings. Future studies will be required to determine how these pathways interact with systemic factors such as inflammation and neural signalling, and whether targeting downstream effectors of contractility, such as ROCK signalling, or rescuing NAD^+^ or αKG with dietary interventions, may provide additional therapeutic benefit.

In summary, we identify a metabolic–epigenetic–mechanical axis linking tumour-derived lactate to skeletal muscle atrophy, through alterations in redox balance, chromatin regulation, and cytoskeletal dynamics. Our work highlighted the importance of metabolism in tumour–muscle crosstalk and highlights new opportunities for therapeutic intervention in cancer cachexia.

## ACKNOWLEDGEMENTS

We thank Julian Downward and Christopher Moore for breeding and providing Kras^G12D^;Lkb1^fl^ (KL) mice, Fabio Zani for generating the KFC cell lines and Marc Hennequart the KVAW12.1g cell line. We thank the Francis Crick Institute’s Advanced Light Microscopy STP and especially Kurt Anderson and Matt Renshaw for microscopy support as well as the following STPs: Metabolomics, Flow Cytometry, Cell Science and Experimental Histopathology. We thank the Francis Crick Institute’s Biological Research Facility (BRF) for their generous support with *in vivo* experiments. The purchase of NL5 microscope was supported by a MRC EQUIP - World Class Labs Award 2024/25 - MC_PC_APP25878 to the Advanced Light Microscopy STP. We are also grateful to the Cancer Grand Challenge CANCAN consortium for helpful discussion and advice. K.H.V. is funded by ERC-2020-ADG PancObese 10102064 and supported by the Francis Crick Institute which receives its core funding from Cancer Research UK (CC2073), the UK Medical Research Council (CC2073) and the Wellcome Trust (CC2073). FST acknowledges funding by The European Union (ERC project no. 759108 – HISTOID; Horizon Europe projects no. 101080690 – MAGIC and no. 101080229 – DREAMS; Views and opinions expressed are however those of the authors only and do not necessarily reflect those of the European Union or HADEA; Neither the European Union nor HADEA can be held responsible for them), Great Ormond Street Hospital Children’s Charity and the Myotubular Trust (V2406), Muscular Dystrophy UK & LifeArc (CMCK-UK.FID119430465) AFM-Telethon and RYR1-Foundation.

## AUTHOR CONTRIBUTIONS

V.P. and K.H.V. designed research; V.P. performed research; F.S.T. advised on the myotube studies,L.R. and V.M.L. provided 2D and 3D samples of immortalised human myoblasts, I.F. performed LC-MS/GC-MS, V.P. and K.H.V. analysed data; and VP and K.H.V. wrote the paper.

## DECLARATION OF INTERESTS

FST and his research group are part of the Horizon Europe “MAGIC” (www.magic-horizon.eu) and “DREAMS” consortia (www.dreamshorizon.eu) which include private SME partners. FST is inventor on a patent application on muscle tissue cultures (WO/2024/261261); he is academic PI of a Crick-AstraZeneca partnership project, and his unrelated consultancies were managed by UCL Consultants. K.H.V. is on the board of directors and shareholder of Bristol Myers Squibb and on the science advisory board (with stock options) of RAZE Therapeutics, Volastra Pharmaceuticals and Kovina Therapeutics. She is on the SAB of Ludwig Cancer, a co-founder and consultant of Faeth Therapeutics and on the Advisory Board of Cell, Cancer Cell, Molecular Cell, Cell Press Blue and Cell Metabolism. She has been in receipt of research funding from Astex Pharmaceuticals and AstraZeneca and contributed to CRUK Cancer Research Technology filing of patent application WO/2017/144877.

The other authors declare no competing interests.

## METHODS

### Cell culture

Mouse KPC or KFC PDAC cell lines were derived from pancreata of individual C57Bl/6 mice with tumours on the pancreas driven by *Pdx1-cre*;LSL-Kras^G12D/+^;LSL-Trp53^R172H/+^ (FC1245, originally isolated by Prof. David Tuveson’s lab, KPC1), *p48-cre*;LSL-Kras^G12D/+^;LSL-Trp53^R172H/+^ (KVAW12.1g, KPC2), *Pdx1-cre*;LSL-Kras^G12D/+^;LSL-Trp53^R172H/+^; Rosa26^YFP/YFP^ (6419c5^69^, KPC3), or *p48-cre*;LSL-Kras^G12D^;LSL-Trp53^R172H^ (KVAT7.3c-KFC1, KVAT8.2c-KFC2, KVAT9.2c-KFC3, KVAT2.1h-KFC4) mice.

Quality control of cell lines was performed by the Francis Crick Institute Cell Science Technology Platform, including mycoplasma testing, STR profiling and species validation. Cells were cultured at 37°C in humidified atmosphere of 5% CO_2_. C2C12, KPC1-3, KFC1-4, PANC1 and MIAPaCa-2 cells were cultured in DMEM (ThermoFisher Scientific, Gibco, 41966029) supplemented with 10% FBS (ThermoFisher Scientific, Gibco, A5256701) and penicillin-streptomycin (10,000 units mL^-1^, ThermoFisher Scientific, Gibco, 15140122). AsPC1 cells were cultured in RPMI 1640 (ThermoFisher Scientific, Gibco, 21875034) supplemented with 10% FBS (ThermoFisher Scientific, Gibco, A5256701) and penicillin-streptomycin (10,000 units mL^-1^, ThermoFisher Scientific, Gibco, 15140122). AB1190 cells were cultured in skeletal muscle growth medium (PromoCell, C23060).

#### C2C12 differentiation

C2C12 cells were plated on fibronectin-coated (Merck, F1141; 10 μg mL^-1^) plastic dishes or collagen-I-coated (Corning, 354236; 100 μg mL^-1^) glass coverslips. Once they reached confluency, cells were washed with PBS and differentiation medium was added consisting of DMEM (ThermoFisher Scientific, Gibco, 21969035) supplemented with Glutamax (ThermoFisher Scientific, 35050038; 2 mM), 1x Insulin-Transferrin-Selenium-Ethanolamine (ITS-X) (ThermoFisher Scientific, 51500056) and penicillin-streptomycin (10,000 units mL^-1^, ThermoFisher Scientific, Gibco, 15140122). Differentiation medium was renewed after 48 hours and cells have differentiated to myotubes typically 72 hours post-induction of differentiation.

#### Immortalised human myoblasts differentiation in 2D

Human immortalised myoblasts (AB1190) were kindly provided by the Myoline platform of the Institute de Myologie, Paris, France. Myoblasts were expanded in Skeletal Muscle Cell Growth Medium (PromoCell, C-23060) at 37°C with 5% CO_2_. Cells were enzymatically detached using TrypLE express (ThermFisher, 12604013) and passaged when reaching 70% confluency. To induce differentiation, cells were plated on Matrigel-coated (Corning, 354234; 1:100) glass-bottom 24 well plates (CellVis, P24-1.5H-N). Once they reached 100% confluency, differentiation to myotubes was induced with a differentiation medium composed of Dulbecco’s Modified Eagle’s Medium – High Glucose (DMEM-HG; 11965092, Gibco) supplemented with Glutamax (Gibco, 35050061; 2 mM) and insulin solution from bovine pancreas (Sigma-Aldrich, I0516; 10 μg mL^-1^). Differentiated myotubes were treated after 2 days with cancer-cell derived metabolites for 24 hours.

#### Human 3D engineered muscle tissues

3D muscle constructs were generated with human immortalised myoblasts as previously described^33^. Briefly, cells were treated with ROCK inhibitor Y27362 (Selleckchem, SMO2-5; 3.3 μg mL^-1^) for 60 min at 37 °C with 5% CO2 and 18% O2. Cells were trypsinized and, for each 3D muscle construct, 10^^6^ cells were mixed on ice with a solution of heat inactivated growth medium (PromoCell, C-23060), Matrigel Growth Factor Reduced (Corning, 356230; 1:10), Fibrinogen (Sigma-Aldrich, cat. no. F8630; 115.2 μg mL^-1^) and thrombin (Sigma-Aldrich, cat. no. T7326; 50 IU). The solution was transferred into 2% agarose molds (UltraPure agarose, Thermo Fisher, cat. no. 16500100), within which were positioned PDMS posts ((EHT Technologies) in 24 well plates, and incubated at 37 °C with 5% CO2 and 18% O2 for 2 h to allow hydrogel polymerisation. Then, warm DMEM-HG was added to the wells to facilitate detachment of the polymerized hydrogel from the agarose molds. After 10 minutes, the PDMS post racks together with the hydrogels were transferred to a new 24-well plate containing myoblast growth medium (PromoCell, C-23060) supplemented with aprotinin (Merck, A3428; 33 μg mL^-1^) and incubated at 37 °C with 5% CO2 and 18% O2 for 48 hours. Then the medium was switched to differentiation medium consisting of DMEM-HG (DMEM-HG; 11965092, Gibco) supplemented with Glutamax (Gibco, 35050061; 2 mM), insulin solution from bovine pancreas (Sigma-Aldrich, I0516; 10 μg mL^-1^) and aprotinin (Merck, A3428; 33 μg mL^-1^). Media was changed every 48 hours. 3D muscle constructs were exposed to conditioned media starting from day 7 post-differentiation.

#### Generation of conditioned media

Mouse KPC, KFC and human (PANC1, MiaPaCa2, AsPC1) pancreatic cancer cells were plated on 15 cm^2^ tissue culture plates to reach confluency 24 hours post plating. Confluent plates were washed twice with PBS and ‘pre-conditioned’ media was added at 10 mL per plate, consisting of DMEM (ThermoFisher Scientific, Gibco, 21969035) supplemented with Glutamax (ThermoFisher Scientific, 35050038; 2 mM), penicillin-streptomycin (10,000 units mL^-1^, ThermoFisher Scientific, Gibco, 15140122) and 1% dialysed FBS (dFBS) (ThermoFisher Scientific, Gibco, 26400044). After 24 hours conditioned media were collected and centrifuged at 300xg for 5 min at 4°C. The supernatant was collected and centrifuged again at 4,000 x g for 10 min at 4°C. The supernatant was then filtered through a 0.22 μm PVDF filter (Merck, SLGVR33RS) before further processing or storage at −70°C.

Conditioned media were also generated as described above with glucose or galactose as the carbon donor. The media composition were DMEM, no glucose, no glutamine, no phenol red (ThermoFisher Scientific, A1443001) supplemented with Glutamax (ThermoFisher Scientific, 35050038; 2 mM), sodium pyruvate (Merck, P2256; 1 mM), 1% dFBS (ThermoFisher Scientific, Gibco, 26400044), penicillin-streptomycin (10,000 units mL^-1^, ThermoFisher Scientific, Gibco, 15140122) and either 12 mM glucose (ThermoFisher Scientific, A2494001) or 12 mM of galactose (Merck, G5388-100G).

#### Fractionation of conditioned media

Conditioned media were either boiled at 95°C to denature protein components or filtered through a Amicon® Ultra Centrifugal Filter, 3 kDa MWCO (Merck; UFC900308) column with 5 mL of media per column centrifuged at 4,000 x g for 60 min at 4°C. From this filtration the concentrated resulting protein fraction (approximately 250 μL) was resuspended to 5 mL final volume with ‘pre-conditioned’ media, whereas the <3 kDa flow-through (metabolite fraction) was used directly. Conditioned media were also fractionated with a Folch’s extraction method; 400 μL of media were added to 500 μL Methanol (Optima LC/MS grade, Fisher Chemical, 10636545) and 1 mL of chloroform (HPLC grade, Merck, 366927) in 2 mL Eppendorf tubes (Eppendorf, 0030120094). The solution was vortexed for 10 sec, incubated on ice for 5 min and then centrifuged at 18,000 x g for 10 min at 4°C. The aqueous phase containing polar metabolites was collected, dried using centrifugal evaporation under vacuum (SpeedVac, RVC 2-33 Cdplus, Martin Christ Gefriertrocknungsanlagen GmbH) and then resuspended with water (HyClone™ HyPure Water, Cell Culture Grade, Cytiva, SH30529.02) at a final volume of 400 μL and incubated at 37°C for 10 min in a Eppendorf Thermomixer. The organic fraction containing lipids was collected into glass vials, dried using centrifugal evaporation under vacuum (SpeedVac, RVC 2-33 Cdplus, Martin Christ Gefriertrocknungsanlagen GmbH) and resuspended with 20 μL ethanol under rotation for 10 min at room temperature and further resuspended to a final volume of 400 μL with ‘pre-conditioned’ media.

#### Treatment of myotubes

Myotubes were treated with complete ‘pre-conditioned’ medium or cancer cell conditioned medium or their derived fractions as indicated mixed at a 50% dilution with myotube maintenance medium ensuring that the essential components ITS-X (C2C12), insulin and/or aprotinin (AB1190, 2D and 3D) were at a 1x final dilution. Myotubes were typically analysed 24 hours post-treatment. Myotubes were also exposed to the following reagents mixed directly with their maintenance medium for 24 hours: sodium-L-lactate (Merck, L7022; 0.4-10 mM), dimethyl 2-oxoglutarate (cell-permeable αKG, Merck, 349631; 0.5 mM), N-acetyl-L-cysteine (NAC, Merck A9165; 5 mM), Glutathione ethyl ester (GSH-EE, cell-permeable glutathione, Cayman Chemical, 14953; 2.5 mM), (±)-6-Hydroxy-2,5,7,8-tetramethylchromane-2-carboxylic acid (Trolox, Merck, 23813; 100 μM), MitoTEMPO (Merck, SML0737; 100 μM), Ferrostatin-1 (Merck, SML0583-5MG; 1 μM), Liproxstatin-1 (Merck, SML1414-5MG; 500 nM), Nicotinamide (NAM, Merck, N0636; 5 mM), β-nicotinamide mononucleotide (βNMN, Merck, N3501; 500 μM), GSK-J4 (Cambridge Bioscience, HY-15648B-10MG; 1 μM).

### Immunofluorescence

#### 2D cultures

Cells were washed with PBS and fixed with 4% formaldehyde for 10 min. Then, they were permeabilised with 0.1% Triton X-100 for 5 min and incubated for 60 min with the following primary antibodies: anti-Myosin Heavy Chain (1/300; mouse monoclonal MF20, Novus Biologicals; Biotechne, MAB4470), mCherry (1/1000; chicken polyclonal, Abcam, ab205402), anti-trimethyl-Histone H3 (Lys27) [H3K27me3] (1/500; rabbit polyclonal, Merck (Millipore), 07-449) and phospho-Myosin Light Chain 2 (Ser19) pMLC2^Ser^^19^ (1/200; rabbit polyclonal, Cell Signaling Technology, 3671).

Detection was performed using the following secondary antibodies: Alexa Fluor 488 donkey anti-rabbit (1/500; Invitrogen, A21206), Alexa Fluor 594 donkey anti-mouse (1/500; Invitrogen, A21203), Alexa Fluor 594 goat anti-chicken (1/500; Invitrogen, A11042), Alexa Fluor 488 goat anti-mouse IgG2b (γ2b) (1/500; Invitrogen, A21141), Alexa Fluor 546 goat anti-mouse IgG1 (γ1) (1/500; Invitrogen, A21123). Nuclei were visualised with DAPI (1 μg mL^-1^ ThermoFisher Scientific, #62248) and F-actin with Alexa Fluor 647 Phalloidin (1/100; ThermoFisher Scientific, A22287).

#### 3D cultures

3D engineered muscle tissues were immunostained as previously described^33^. The hydrogels were fixed in 4% PFA at 4°C for 4 hours, removed from their PDMS posts and transferred to a well containing blocking buffer consisting of 10% FBS, 0.5% Triton X-100 and 1% BSA in Tris-buffered saline (TBS) at pH 7.5. After 6 hours incubation at 4°C, the following antibodies were added in permeabilization buffer (0.5% Triton X-100 and 1% BSA in TBS pH 7.4) at 4°C for 16 hours: anti-Myosin Heavy Chain (1/300; mouse monoclonal MF20, Novus Biologicals; Biotechne, MAB4470) and anti-trimethyl-Histone H3 (Lys27) [H3K27me3] (1/500; rabbit polyclonal, Merck (Millipore). Then, six washes for 60 min each were performed in TBS1x at RT on a plate shaker. The following secondary antibodies were added in permeabilization buffer (0.5% Triton X-100 and 1% BSA in TBS pH 7.4) at 4°C for 16 hours: Alexa Fluor 488 donkey anti-rabbit (1/500; Invitrogen, A21206), Alexa Fluor 594 donkey anti-mouse (1/500; Invitrogen, A21203). Nuclei were visualised with DAPI (1 μg mL^-1^ ThermoFisher Scientific, #62248) and hydrogels were mounted in concave coverslips with ProLong™ Diamond Antifade mountant (ThermoFisher Scientific, P36961).

#### Imaging

Images were acquired using a Zeiss AxioImager.M1 microscope equipped with a Zeiss Plan-Neofluar 20x 0.5 objective or a Zeiss Observer.Z1 microscope equipped with a Zeiss Plan-Neofluar 10x 0.3 objective and using MicroManager (version 2.0)^70^ software for acquisition. High resolution images were acquired with a Zeiss Invert880 microscope equipped with a 63x/1.4 Plan Apochromat objective or a Nikon SoRa spinning disk with a 40x CFI Apo LWD λS water immersion objective. All images were processed with Fiji software (ImageJ v2.16.0).

### Live imaging

Differentiated C2C12 myotubes on 8-well glass-bottom chambers (C8-1.5H-N) were exposed to control or cancer cell conditioned media containing the cell permeable actin probe SiR-actin (Tebubio, Spirochrome, 251SC001; 1/2000) and the nuclei marker SPY-505 DNA (Tebubio, Spirochrome, SC101; 1/1000). Cells were imaged within a contained unit at 37 °C and under 5% CO2 using a NL5+ line rescanning confocal (Confocal.NL) on an IX83 inverted microscope (Olympus), 60x/1.3 NA UPlan S Apo objective and Orca Quest 2 camera (Hamamatsu), controlled via Micro-Manager v2.0 software with images being acquired at 20 min intervals for 16 hours. Environmental conditions maintained with a Boldline 3 environmental chamber (Okolab). All images were processed with Fiji software (ImageJ v2.16.0) and myotube nuclei were tracked using the MTrackJ plugin (ImageJ).

### Western blotting

Proteins were extracted in RIPA Lysis and Extraction buffer (ThermoFisher Scientific, 89900) supplemented with protease (cOmplete, EDTA-free Protease Inhibitor Cocktail, Roche, Merck, 4693159001) and phosphatase (PhosSTOP, Roche, Merck, 4906845001) inhibitors. Lysates were cleared by centrifugation at 18,000 x g for 10 min at 4°C. Total protein content was quantified by Pierce BCA assay (ThermoFisher Scientific, 23227). Following normalisation to total protein content and addition of NuPAGE LDS Sample Buffer (4x) (ThermoFisher Scientific, NP0007) supplemented with NuPAGE™ Sample Reducing Agent (ThermoFisher Scientific NP0004), lysates were heated to 95°C for 5 min or 37°C for 20 min (Oxphos cocktail) and typically 20 μg of proteins were resolved in NuPAGE 4-12% bis-tris polyacrylamide gels (Thermo Fisher Scientific, NP0336BOX) with NuPAGE MOPS SDS Running Buffer (Thermo Fisher Scientific, NP000102) or NuPAGE™ MES SDS Running Buffer (Thermo Fisher Scientific, NP000202). Proteins were transferred to nitrocellulose (ThermoFisher Scientific, IB23001) or PVDF membranes (ThermoFisher Scientific, IB24001) using semi-dry transfer (ThermoFisher Scientific, IB21001). Membranes were blocked for 60 min using 5% milk and primary antibodies against FBXO32 (1/1000; mouse monoclonal EPR9148(2), Abcam, ab168372), Myogenin (MyoG, 1/1000; mouse monoclonal F5D, ThermoFisher Scientific, 14-5643-82), pMLC2^Ser19^ (1/1000; rabbit polyclonal, Cell Signaling Technology, 3671), MLC2 (1/1000; rabbit polyclonal, Cell Signaling Technology, 3672), the total oxphos rodent western blot antibody cocktail (6 μg mL^-1^; mouse, Abcam, ab110413) and Vinculin (1/2000; mouse monoclonal 7F9, Santa Cruz Biotechnology, sc-73614) were incubated overnight at 4 °C in blocking solution. Membranes were washed three times in TBS + 0.2% TWEEN-20 (Merck, P7949) and incubated with secondary antibodies: goat anti-rabbit IgG DyLight^TM^ 800nm (ThermoFisher Scientific, SA5-35571), donkey anti-mouse IgG AlexaFluor^TM^ 680 (ThermoFisher Scientific, A10038) for 60 min at room temperature and washed 3 times in TBS + 0.2% TWEEN-20. Fluorescence signal was captured using a LI-COR Odyssey CLx Imaging System (LI-COR Biosciences) with Image Studio software (version 5.2). For total oxphos cocktail a horse anti-mouse IgG HRP-linked antibody was used (Cell Signaling Technology, 7076). Membranes were washed 3 times in TBS + 0.2% TWEEN-20 and developed with Pierce ECL chemiluminescence kit (Thermo Fisher Scientific, 32106) using a ChemiDoc MP imaging system (Bio-Rad).

### Quantification of gene expression

RNA was isolated from 6-well plates using the Monarch Spin RNA Isolation kit (NEB, T2110S) according to the manufacturer’s instructions. For RNA from tissue, approximately 50 mg of tissue were homogenized in 1 mL of TRIzol (ThermoFisher Scientific, 15596026) using Precellys ceramic kit 1.4 mm 2 mL tubes (VWR, 432-3751) and a Precellys 24 homogenizer (Bertin Technologies). Following addition of 200 μL chloroform, the solution was centrifuged for 10 min at 18,000 x g and 4 °C. The aqueous layer was mixed with isopropanol (50%) and centrifuged for 15 min at 18,000 x g and 4 °C. The pellet was washed with 75% ethanol and collected by centrifugation for 5 min at 7,500 x g and 4 °C. Following resuspension with 30 μL of nuclease-free water, the RNA was DNase treated using the Qiagen RNase-Free DNase set (Qiagen, 79254) and cleaned with the RNeasy Mini kit (Qiagen, 74104) according to manufacturer’s instructions. Measurements of RNA concentration and purity were routinely performed using a DeNovix DS-11FX+ spectrophotometer before downstream processing. cDNA synthesis was performed with the LunaScript RT SuperMix kit (NEB, E3010L) using typically 0.5 μg of RNA. Quantitative PCR with reverse transcription (qRT-PCR) analysis was performed in technical triplicates using QuantStudio™ 7 Flex system (Applied Biosystems, Thermo Fisher Scientific) with the Luna Universal qPCR Master Mix (NEB, M3003X). Relative mRNA quantification was performed using the 2^-ddCT method for multiple genes. A list of the qRT-PCR primer sequences is provided in Supplementary Table 1.

### Cloning

The plasmids pUC57-LbNOX (Addgene #75285; http://n2t.net/addgene:75285; RRID:Addgene_75285) and pUC57-mitoLbNOX (Addgene #74448; http://n2t.net/addgene:74448; RRID:Addgene_74448) were a gift from Vamsi Mootha^48^. The ORFs of LbNOX and mitoLbNOX were PCR amplified and cloned upstream of a T2A-mCherry encoding sequence into the pcDNA3.1(+) vector using the Gibson assembly kit (NEB, E5510S) according to the manufacturer’s instructions. DNA construct information is provided in Supplementary Table 1.

### Transfection of DNA constructs

Typically, 2×10^4^ cells per well were plated on 96-well plates. After 16 hr, 1 μg of plasmid DNA was mixed with 5 μL of Lipofectamine 2000 transfection reagent (Thermo Fisher Scientific, 11668019) in serum-free medium to a total volume of 200 μL. Following incubation at room temperature for 5 min, DNA/Lipofectamine 2000 mix was added to cells. 4 hours post-transfection, media were switched to differentiation media which were renewed 48 hours afterwards. Treatment with conditioned media started 72 hours post transfection and cells were fixed and analysed after 24 hours.

### Metabolite extraction from media

Typically, 10 μL of medium per replicate was lysed in ice-cold extraction buffer consisting of 50 μL of chloroform (HPLC grade, Merck, 366927), 150 μL methanol (Optima LC/MS grade, Fisher Chemical, 10636545) and 140 μL water (OPTIMA, LC/MS Grade, Fisher Scientific, 10505904) containing 1 nmol *scyllo*-inositol (Merck, I8132) and 1 nmol L-Norleucine (Merck, N8513). All lysates were vortexed and then centrifuged at 18,000 x g for 10 minutes at 4 °C. The aqueous phase was transferred to a fresh Eppendorf extraction tube, dried using centrifugal evaporation under vacuum (SpeedVac, RVC 2-33 Cdplus, Martin Christ Gefriertrocknungsanlagen GmbH) and analysed using GC/MS.

### Calculation of metabolite exchange rates

For each identified polar metabolite, total fmol per well detected in media at t = 24 hours were subtracted from the total fmol per well detected in media at t = 0 hours and divided by corresponding cell numbers, providing fmol per cell consumed/released over a period of 24 hours.

### [U-^13^C_6_]D-glucose tracing

Differentiated myotubes were treated for 16 hours with 50% myotube maintenance medium [DMEM, no glucose, no glutamine, no phenol red (ThermoFisher Scientific, A1443001) supplemented with Glutamax (ThermoFisher Scientific, 35050038; 2 mM), sodium pyruvate (Merck, P2256; 1 mM), penicillin-streptomycin (10,000 units mL^-1^, ThermoFisher Scientific, Gibco, 15140122), 2x ITS-X (ThermoFisher Scientific, 51500056) and 12 mM glucose (ThermoFisher Scientific, A2494001)] and 50% metabolites-derived from control or cancer cell conditioned media generated with DMEM, no glucose, no glutamine, no phenol red (ThermoFisher Scientific, A1443001) supplemented with Glutamax (ThermoFisher Scientific, 35050038; 2 mM), sodium pyruvate (Merck, P2256; 1 mM), 1% dFBS (ThermoFisher Scientific, Gibco, 26400044), penicillin-streptomycin (10,000 units mL^-1^, ThermoFisher Scientific, Gibco, 15140122) and 12 mM glucose (ThermoFisher Scientific, A2494001). Then the medium was replaced with DMEM, no glucose, no glutamine, no phenol red (ThermoFisher Scientific, A1443001) containing 12 mM [U-^13^C_6_]D-glucose, 99% (Merck, 389374) and Glutamax (ThermoFisher Scientific, 35050038; 2 mM), sodium pyruvate (Merck, P2256; 1 mM), penicillin-streptomycin (10,000 units mL^-1^, ThermoFisher Scientific, Gibco, 15140122), 50% conditioned media derived metabolites and 1x ITS-X (ThermoFisher Scientific, 51500056). After 4 hours, cells were washed three times with ice-cold PBS and metabolites were extracted with ice-cold extraction buffer, consisting of 200 μL chloroform (HPLC grade, Merck, 366927), 600 μL methanol (Optima LC/MS grade, Fisher Chemical, 10636545) and 200 μL water (OPTIMA, LC/MS Grade, Fisher Scientific, 10505904). Two independent wells per condition were pooled per replicate. Samples were centrifuged at 18,000 x g for 10 minutes at 4 °C and the supernatant was transferred to a fresh Eppendorf extraction tube and dried using centrifugal evaporation under vacuum (SpeedVac, RVC 2-33 Cdplus, Martin Christ Gefriertrocknungsanlagen GmbH). Samples were resuspended in 50 µL chloroform (HPLC grade, Merck, 366927), 150 µL methanol (Optima LC/MS grade, Fisher Chemical, 10636545), and 150 µL water (OPTIMA, LC/MS Grade, Fisher Scientific, 10505904) containing 1 nmol *scyllo*-inositol (Merck, I8132) and 1 nmol L-Norleucine (Merck, N8513). The aqueous phase was transferred to a fresh Eppendorf extraction tube, dried using centrifugal evaporation under vacuum (SpeedVac, RVC 2-33 Cdplus, Martin Christ Gefriertrocknungsanlagen GmbH) and analysed using GC/MS. Metabolite values were normalized to corresponding cell numbers per condition to calculate nmol per cell.

### [U-^13^C_5_]L-glutamine tracing

Differentiated myotubes were treated for 24 hours with 50% myotube maintenance medium [DMEM, no glucose, no glutamine, no phenol red (ThermoFisher Scientific, A1443001) supplemented with sodium pyruvate (Merck, P2256; 1 mM), penicillin-streptomycin (10,000 units mL^-1^, ThermoFisher Scientific, Gibco, 15140122), 2x ITS-X (ThermoFisher Scientific, 51500056) and 12 mM glucose (ThermoFisher Scientific, A2494001)] and 50% metabolites-derived from control or cancer cell conditioned media generated with DMEM, no glucose, no glutamine, no phenol red (ThermoFisher Scientific, A1443001) supplemented with Glutamine (ThermoFisher Scientific, 35050038; 2 mM), sodium pyruvate (Merck, P2256; 1 mM), 1% dFBS (ThermoFisher Scientific, Gibco, 26400044), penicillin-streptomycin (10,000 units mL^-1^, ThermoFisher Scientific, Gibco, 15140122) and 12 mM glucose (ThermoFisher Scientific, A2494001). The media also contained 2 mM [U-^13^C_3_]L-glutamine (Merck, 605166). After 24 hours, metabolites were extracted with ice-cold extraction buffer, consisting of methanol (Optima LC/MS grade, Fisher Chemical, 10636545) / acetonitrile (Optima LC/MS grade, Fisher Chemical, 10055454) / H2O (OPTIMA, LC/MS Grade, Fisher Scientific, 10505904) (50:30:20). Lysis buffer volume added per well was adjusted to cell number, using cell-count numbers from matching plates. Samples were centrifuged at 18,000 x g for 10 minutes at 4 °C and the supernatant was transferred to a fresh Eppendorf extraction tube and dried using centrifugal evaporation under vacuum (SpeedVac, RVC 2-33 Cdplus, Martin Christ Gefriertrocknungsanlagen GmbH). Samples were then resuspended to 70 μL of acetonitrile (Optima LC/MS grade, Fisher Chemical, 10055454) / H2O (OPTIMA, LC/MS Grade, Fisher Scientific, 10505904) (80:20). Samples were analysed using LC/MS.

### Amino acid quantification in muscle tissue and serum

Tissue samples were snap-frozen in liquid nitrogen and stored at −80 °C. Samples were homogenized in a pre-chilled (−20 °C) solvent mixture consisting of methanol, acetonitrile, and water (50:30:20, v/v), all of LC/MS-grade quality (Optima, Fisher Scientific). 1 mL of solvent was used per 50 mg of tissue and samples were homogenised using a Precellys ceramic kit 1.4 mm 2 mL tubes (VWR, 432-3751) and a Precellys 24 homogenizer. Lysates were incubated on ice for 60 min and then centrifuged at 18,000 x g for 10 min at 4°C. Supernatants were collected to a fresh Eppendorf extraction tube and dried using centrifugal evaporation under vacuum (SpeedVac, RVC 2-33 Cdplus, Martin Christ Gefriertrocknungsanlagen GmbH). Samples were then resuspended to 100 μL of acetonitrile (Optima LC/MS grade, Fisher Chemical, 10055454) / H2O (OPTIMA, LC/MS Grade, Fisher Scientific, 10505904) (80:20). Samples were analysed using LCMS.

Blood was isolated with cardiac puncture after S1K culling in Heparin-treated tubes. Following incubation at room temperature for > 20 min, plasma was isolated by centrifugation at 3000 x g for 15 min at 4°C and was stored at −70 °C. 10 μL of serum was extracted with 30 μL of Methanol (Optima LC/MS grade, Fisher Chemical, 10636545) for 5 min on ice and centrifuged for 10 min at 4 °C and 18,000 x g. Samples were dried using centrifugal evaporation under vacuum (SpeedVac, RVC 2-33 Cdplus, Martin Christ Gefriertrocknungsanlagen GmbH) and partitioned in 50 μL chloroform, 150 μL methanol and 150 μL H2O buffer. Samples were centrifuged for 10 min at 4 °C and 18,000 x g. The polar phase was then dried using SpeedVac and was resuspended in 100 μL acetonitrile (Optima LC/MS grade, Fisher Chemical, 10055454) / H_2_O (Optima, LC/MS Grade, Fisher Scientific, 10505904) (80:20, v/v) buffer. Samples were analysed using LCMS.

### Liquid chromatography-mass spectrometry (LC-MS)

Metabolite analysis was performed by LCMS using a Q-EXACTIVE Plus (Orbitrap) mass spectrometer (ThermoFisher) coupled with a Vanquish UHPLC system (ThermoFisher). The chromatographic separation was performed on Atlantis Premier BEH z-HILIC column (100 × 2.1 mm, 1.7 µm; Waters, Milford, MA, USA) was used. Solvent A consisted of 12 mmol L⁻¹ ammonium carbonate (pH 9.0), and solvent B was ACN. The column temperature was maintained at 25 °C, with a flow rate of 0.3 mL min^-1^ and an injection volume of 5 µL. The elution gradient was as follows: 95 % solvent B was held for 1 minute, then decreased linearly to 49% over 10 minutes (1–11 min), followed by a 2-minute hold (11– 13 min). From 13.01 to 15.00 minutes, solvent B was further decreased to 25 %. At 15.01 minutes, it was returned to 95 % and held until 21 minutes. MS was performed with positive/negative polarity switching using a Q-Exactive Plus Orbitrap (ThermoFisher) with a HESI II probe. MS parameters were as follows: spray voltage 3.5 and 3.2 kV for positive and negative modes, respectively; probe temperature 320°C; sheath and auxiliary gases were 30 and 5 arbitrary units, respectively; and full scan range: 65–975 m/z with settings of AGC target and resolution as balanced and high (1e6 and 70,000), respectively. Data were recorded using Xcalibur 4.2.47 software (ThermoFisher). Mass calibration was performed for both ESI polarities before analysis using the standard ThermoFisher Calmix solution. To enhance calibration stability, lock-mass correction was also applied to each analytical run using ubiquitous low-mass contaminants. Data dependent acquisition (DDA) parameters were the following: resolution 17,500; resolution 17,500; AGC targeted 2e5; maximum IT 100 ms; Stepped NCE were: 20, 30, 40. Metabolites were identified and quantified by accurate mass and retention time and by comparison to the retention times, mass spectra, and responses of known amounts of authentic standards using TraceFinder 5.1 EFS software (ThermoFisher). Samples have been supplemented with 5 μM [^13^C ^15^N]L-Valine (Cambridge Isotope Laboratories, Inc.; CNLM-442-H-0.25) during the extraction process.

### Gas chromatography-mass spectrometry (GC/MS)

Dried samples were washed three times with 30 μL methanol and derivatised first with 20 μL Methoxyamine hydrochloride (Merck, 89803) solution in pyridine (Merck, 270970) (20 mg mL^-1^) for 16 hours and then with 20 μL BSTFA+1% TMCS silylation reagent (Thermo Fisher Scientific, TS-38831) for 60 min. Metabolite analysis was performed by GC-MS using an Agilent 7890B-7000C system. Splitless injection (injection temperature 250 °C) onto a 30 m + 10 m × 0.25 mm DB-5MS+DG column (Agilent J&W) was used, using helium as the carrier gas, in electron ionization (EI) mode. The initial oven temperature was 70 °C (2 min), followed by temperature gradients to 295 °C at 12.5 °C/min, then to 320 °C at 25 °C/min (held for 3 min). Metabolites were identified by comparison with the retention times and mass spectrum of authentic standards using MassHunter Workstation software (B.06.00 SP01, Agilent Technologies). Abundance was calculated by comparison to responses of known amounts of authentic standards. Data analysis was performed using in house-developed software MANIC (version 3.0), based on the software package GAVIN^71^. Label incorporation was calculated by subtracting the natural abundance of stable isotopes from the observed amounts. Metabolites were identified and quantified in comparison to authentic standards and *scyllo*-Inositol as an internal standard (Sigma, I8132).

### ADP/ATP ratio quantification

ADP/ATP ratio was assessed with a bioluminescent assay (Merck, MAK135) 24 hours post-treatment in 96-well plates according to the manufacturer’s instructions. Briefly, media were removed and cells were washed twice with PBS. 90 μL ATP reagent mix was added to each well. After 1 min incubation at room temperature, luminescence was recorded (RLU_A_). After 10 min incubation at room temperature, luminescence was recorded (RLU_B_) and 5 μL ADP reagent was added per well. After 1 min incubation at room temperature luminescence was recorded (RLU_C_) and ADP/ATP ratio was calculated using the below formula: (RLU_C_-RLU_B_)/ RLU_A_. Luminescence was measured using a Spark multimode microplate reader (Tecan Trading AG).

### NAD^+^/NADH ratio quantification

NAD^+^/NADH ratio was assessed with a colorimetric assay (Abcam, ab65348) 24 hours post-treatment according to the manufacturer’s instructions. Briefly, cells were lysed with 450 μl NADH/NAD^+^ extraction buffer and then centrifuged for 5 min at 4 °C at 20,000 x g. Supernatants were filtered through 10-kDa spin column (Amicon^®^ Ultra Centrifugal Filter, 10 kDa MWCO, Merck, UFC501024) centrifuged at 10,000 x g for 10 min. Then, 200 μL were used for total NAD+ and NADH quantification, while the remaining solution was heated at 60 °C for 30 min to degrade NAD+. 50 μL of each sample were mixed with 100 μL of reaction mixture and incubated for 5 min at room temperature. Following addition of 10 μL developing solution, the reaction was incubated for 1 hour at room temperature and absorbance was measured at 450 nm using a Spark multimode microplate reader (Tecan Trading AG). NAD+ and NADH amounts were calculated using a NADH standard curve. The NAD+/NADH ratio was calculated using the below formula: (NAD(H)total – NADH)/NADH.

### GSH/GSSG ratio quantification

GSH/GSSG ratio was assessed with a bioluminescent assay (Promega, V6611) 24 hours post-treatment according to the manufacturer’s instructions. Briefly, media were removed from 96-well plates and 50 μL per well of either Total Glutathione Lysis Reagent or Oxidized Glutathione Lysis Reagent were added in parallel wells. The plate was incubated for 5 min at room temperature and 50 μL of Luciferin Generation Reagent were added. After 30 min incubation at room temperature 100 μL of Luciferin Detection Reagent were added. Plates were incubated at room temperature for 15 min and luminescence was measured using Spark multimode microplate reader (Tecan Trading AG). The average background luminescence was subtracted from all wells and GSH/GSSG ratio was calculated with the below formula: (Net total glutathione RLU – Net GSSG RLU) / (Net GSSG RLU / 2)

### Plasma Lactate quantification

Lactate was quantified in plasma samples using a colorimetric assay (Abcam, ab65331) according to the manufacturer’s instructions. 5 μL of plasma were mixed with assay buffer, developer solution and enzyme mix and following 30 min incubation at room temperature and the OD 450 nm absorbance was measured with a Spark multimode microplate reader (Tecan Trading AG). Absorbance values were corrected for background signal as well as the blank, and lactate concentration was calculated as follows using a standard curve: ((amount of Lactate in sample calculated from standard curve) / volume of sample added into the well (μL)) x dilution factor.

### Mitochondrial mass and potential quantification

For quantifying mitochondrial mass and mitochondrial membrane potential, cells were labelled using MitoTracker Green (200 nM; M7514, ThermoFisher Scientific) and TMRE-Mitochondrial Membrane Potential Assay Kit (100 nM; ab113852, Abcam) respectively according to the manufacturer’s instructions. MitoTracker Green (200 nM) and TMRE (100 nM) were added for 30 min. Live cells were collected and washed twice with PBS before being analysed with a Fortessa flow cytometer (BD Biosciences). Data were analysed using FlowJo software (v.10.10.0, FlowJo). DAPI (0.5 μg ml^−1^) was used to identify live and dead populations.

### Cellular and mitochondrial ROS quantification

Cellular ROS were quantified using CellROX Deep Red reagent (1 μM; ThermoFisher Scientific, C10422) and mitochondrial ROS using MitoSOX Red (2.5 μM; ThermoFisher Scientific, M36008) according to the manufacturer’s instructions. Briefly, CellROX Deep Red and MitoSOX were added for 20 min and live cells were collected and washed twice with PBS before being analysed with a Fortessa flow cytometer (BD Biosciences). Data were analysed using FlowJo software (v.10.10.0, FlowJo). DAPI (0.5 μg ml^−1^) was used to identify live and dead populations.

### Histology and Immunohistochemistry

Tissues were fixed in 10% neutral buffered formalin for 24 hours. The fixed tissues were processed using Sakura Tissue-Tek VIP® 6 AI Tissue Processor and paraffin embedded. 3μm sections were cut on glass slides. The slides were baked at 60°C for 1 hour and then deparaffinized in xylene and rehydrated using a series of graded industrial methylated spirits solutions and distilled water. For H&E staining the slides were stained using Tissue-Tek Prisma® Plus Automated Slide Stainer. For immunohistochemistry, heat-mediated antigen retrieval was performed using pH 9 Tris-EDTA buffer for 20 minutes in a microwave. Endogenous peroxidase blocking was performed by incubating the slides in Bloxall (SP-6000-100, Vector) for 10 minutes at room temperature. The slides were incubated with 2.5% Normal Horse Serum for 30 minutes at room temperature. Primary antibodies Anti-Slow Skeletal Myosin Heavy chain, 1:1000 (abcam ab234431, LOT 1121984-6) and Anti-Fast Myosin Skeletal Heavy chain antibody, 1:12000 (abcam ab91506, LOT 1039672-4) were applied for 1 hour at room temperature. The slides were incubated in ImmPRESS anti-rabbit Polymer Reagent (Vector MP-7401) for 30 minutes at room temperature. Slides were incubated with 3,3-diaminobenzidine (DAB) chromogen (SK-4105, Vector) for 10 minutes at room temperature. The slides were then counterstained using Harris Haematoxilin, dehydrated, cleared and mounted using Sakura Tissue-Tek Prisma automated stainer. The slides were scanned using Zeiss Axioscan 7 Slide Scanner using a 10X brightfield scanning profile.

### Animals

All *in vivo* experiments were conducted in compliance with the with the Animals (Scientific Procedures) Act 1986 and the EU Directive 2010 (PPL PP1389725) and within institutional welfare and ethical guidelines of the Francis Crick Institute (establishment licence X49D044E1). Mice had ad libitum access to food and water and were kept in a 12h day/night cycle (7:00 to 19:00) in rooms at 21°C and 55% humidity. Mice were acclimatized to their environment for at least one week prior to experimentation. Mice used in the experiments were randomly assigned to experimental groups. C57BL/6 were obtained from the in-house breeding facility.

#### Subcutaneous injections

2×10^5^ cells KPC1 (FC1245) cells were resuspended in 100 μL PBS and introduced into C57BL/6 mice (males, 9-week old; Francis Crick Institute) by subcutaneous injection on the flank. Tumour measurement and body weight monitoring was performed twice weekly by staff blinded to the experimental outcome. After 14 days animals were humanely culled using Schedule 1 methods. Each mouse was an experimental unit and control mice received a 100 μL PBS injection.

#### KL model

The mouse model of Kras^LSL-G12D/+^; Lkb1^f/f^ lung cancer^72^ was in C57BL/6 mice (Francis Crick Institute). Lung tumours were induced in male and female mice, aged 7-10 weeks, using 10^7^ plaque-forming units of Adenovirus CMV-Cre (Ad5CMV-Cre) acquired from the University of Iowa Gene Transfer Vector Core (Iowa City, IA). Body weight was monitored twice per week since Cre administration. Mice were humanely euthanized using a pentobarbital injection, upon exhibiting humane endpoint symptoms.

### Statistics and reproducibility

All datasets were analysed and plotted using Prism 11 (v11.0.0; GraphPad Software) unless otherwise stated. Bubble and spider plots were generated with R Studio (Version 2023.06.1+524). Heatmaps on figures 3 & extended data figure 3 were generated with Metaboanalyst 6.0^73^. Differences between groups were tested for normal distribution and analysed using the appropriate statistical test, as mentioned in each figure legend. Error bars represent s.d., unless otherwise stated. LCMS and GCMS metabolomic measurements of conditioned media and cell lysates were performed within the same day with the indicated number of biological replicates. Biological replicates are cells grown in separate plates/wells in which experimental conditions were replicated within an experiment and each well yielded a sample that was analyzed by LC/MS or GC/MS independently. The remaining experiments were performed independently at least two times and as indicated on figure legends. Biorender.com was used to create illustrations.

### Data and resources availability

Original datasets, analyses and methodological details are available upon request. Information regarding reagents and animal studies can also be found in associated supplementary tables.

---

**Extended Data Figure 1:**
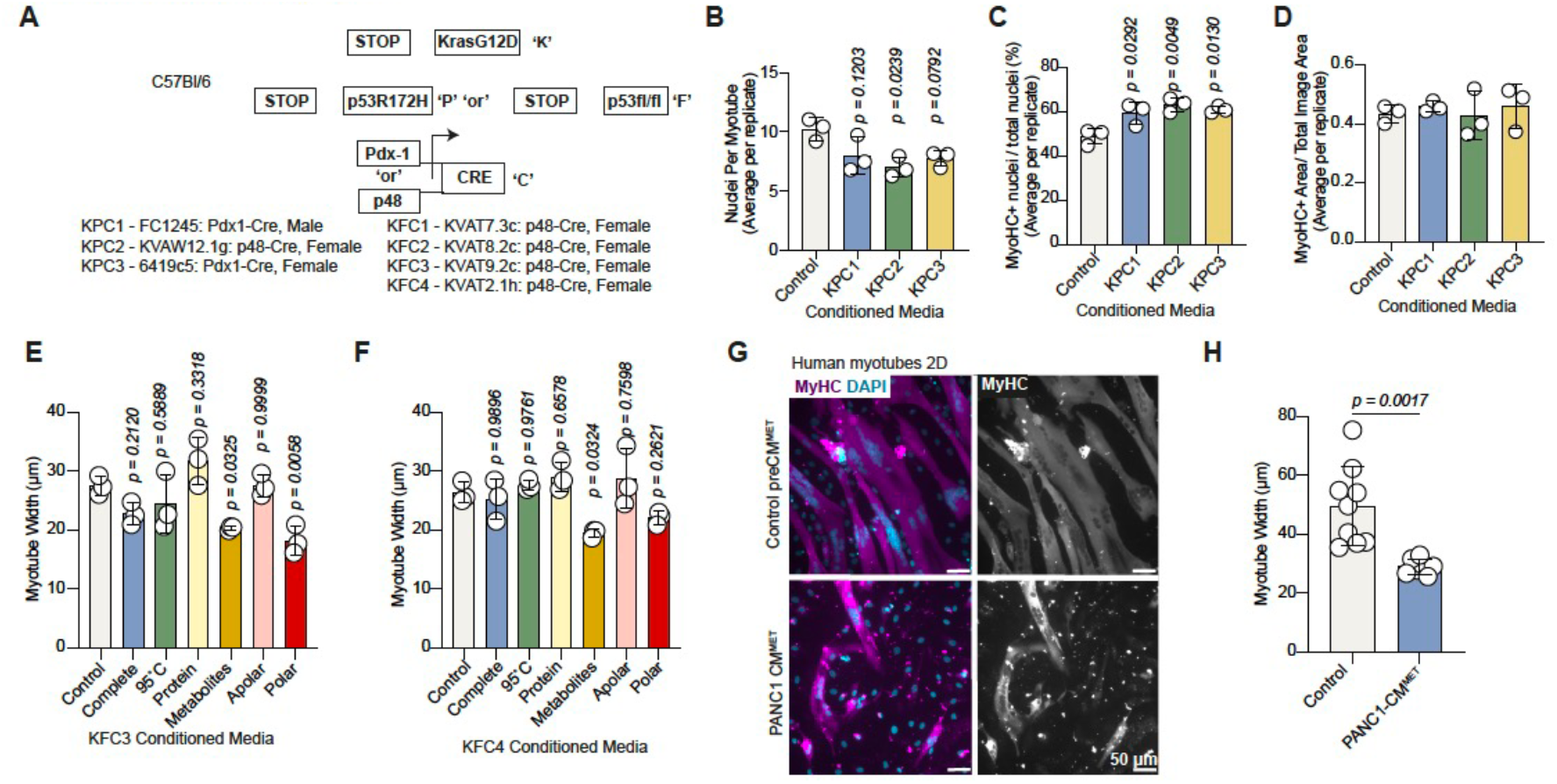
PDAC cell derived metabolites induce myotube atrophy. (A) Schematic indicating the mouse pancreatic cancer cell lines used. (B) Number of nuclei per myotube (fusion index) of myotubes treated with conditioned media as indicated. Values are mean ±s.d. and average of >10 myotubes per replicate. (C) Percentage of MyHC positive nuclei over total number of nuclei per field of view of myotubes treated with conditioned media as indicated. Values are mean ±s.d. and average of >10 myotubes per replicate. (D) Ratio of MyHC positive area over total image area of myotubes treated with conditioned media as indicated. Values are mean ±s.d. and average of >10 myotubes per replicate. (E-F) Width (μm) of myotubes treated with conditioned media fractions from KFC3 **(E)** and KFC4 **(F)**cells as indicated. Values are mean ± s.d. from average of >10 myotubes per replicate. (G) Representative images of MyHC (magenta and grey right panel) and DAPI (blue)-stained human myotubes treated with control or PANC1-derived metabolites. Scale bars; 50 μm. (H) Width (μm) of myotubes from **(G)**. Values are mean ± s.d. from average of >10 myotubes per replicate and from two independent experiments. Statistical significance was assessed by two-tailed Welch’s *t*-test. B, C: Statistical significance was assessed with ordinary one-way analysis of variance (ANOVA) and Tukey’s multiple comparisons test and replicates are from 3 independently produced conditioned media batches. E, F: Statistical significance was assessed with ordinary one-way analysis of variance (ANOVA) and Dunnett’s multiple comparisons test.

**Extended Data Figure 2:**
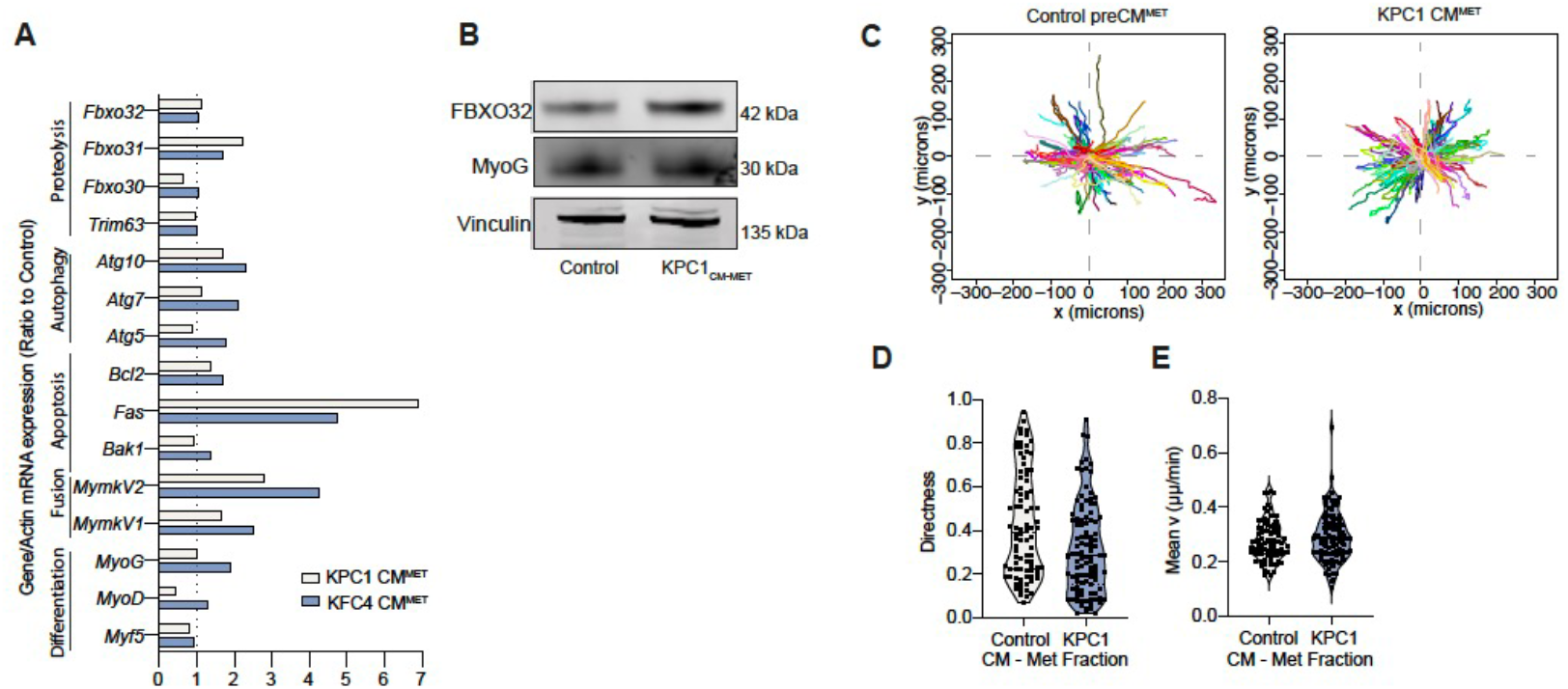
Myotube atrophy induced by PDAC cell-derived metabolites is defined by cytoskeletal stress through increased myosin-dependent contractility. (A) mRNA expression levels of indicated genes in C2C12 myotubes treated with Control, KPC1- or KFC4-derived metabolites. Values are ratio to control and representative from three independent experiments. Ct values were normalised to *Actin*. (B) Immunoblots of FBXO32, MyoG and Vinculin (loading control) in C2C12 myotubes treated with Control or KPC1-derived metabolites. Blots are representative of three independent experiments. (C) Tracks of nuclei movement within myotubes treated with control (left) or KPC1-derived (right) metabolites over 20 hours from treatment start. (D-E) Violin plots of myotube nuclei directness **(D)** and mean velocity (μm/min) **(E)**. Values are mean+s.d. from four replicates per condition and representative from three independent experiments. Data are shown as average per replicate on Fig. 2D & E.

**Extended Data Figure 3:**
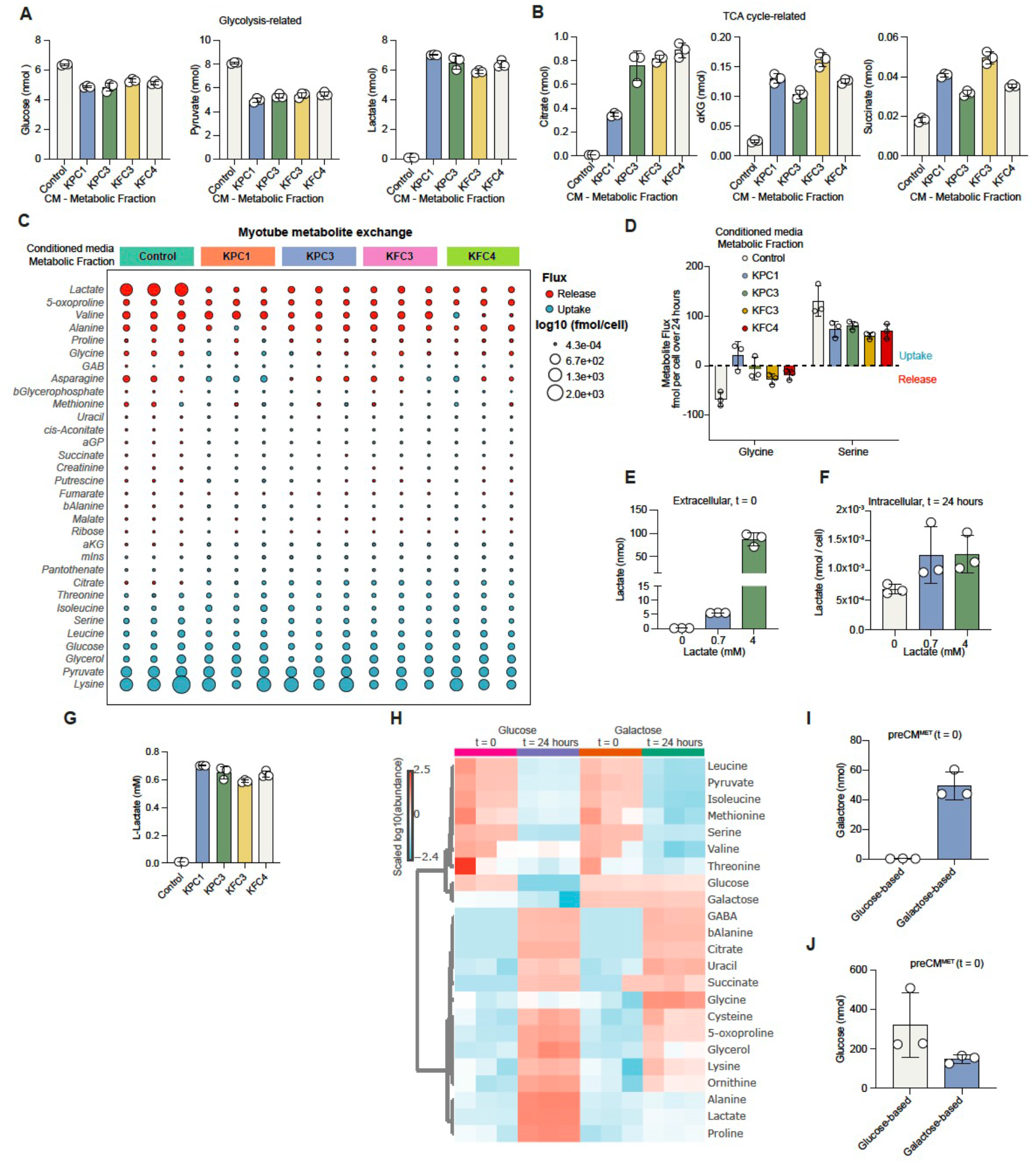
PDAC cell-secreted lactate induces myotube atrophy. (A) Abundance of indicated Glycolysis-related metabolites in metabolite fraction from control or PDAC cell conditioned media. Values are mean±s.d. of three replicates. (B) Abundance of indicated TCA cycle-related metabolites in metabolite fraction from control or PDAC cell conditioned media. Values are mean±s.d. of three replicates. (C) Bubble plot indicating uptake (cyan) or release (red) of indicated metabolites in myotubes treated with control or PDAC cell derived metabolites as indicated for 24 hr. Values are log10 (fmol/cell) from three replicates and bubble size represents the degree of uptake or release. (D) Metabolite flux (uptake or release, fmol per cell) of glycine and serine in myotubes treated with control or PDAC cell derived metabolites as indicated for 24 hr. Values are mean±s.d. from three replicates. (E) Lactate abundance (nmol) in media supplemented with increasing lactate concentration. Values are mean±s.d. from three replicates. (F) Lactate abundance (nmol/cell) in cells supplemented with increasing L-lactate concentration. Values are mean±s.d. from three replicates. (G) L-lactate concentration in metabolite fraction from control or PDAC cell conditioned media. Values are mean±s.d. of three replicates. Concentration values correspond to data shown in (A). (H) Heatmap, abundance of detected metabolites in metabolite fractions of media from glucose or galactose-fed KPC1 cells at t = 0 (preconditioned) and t = 24 hrs (conditioned). Values are scaled log10 (nmol) from three replicates. (I) Galactose abundance (nmol) in galactose and glucose-based media at t = 0. Values are mean±s.d. from three replicates. (J) Glucose abundance (nmol) in galactose and glucose-based media at t = 0. Values are mean±s.d. from three replicates.

**Extended Data Figure 4:**
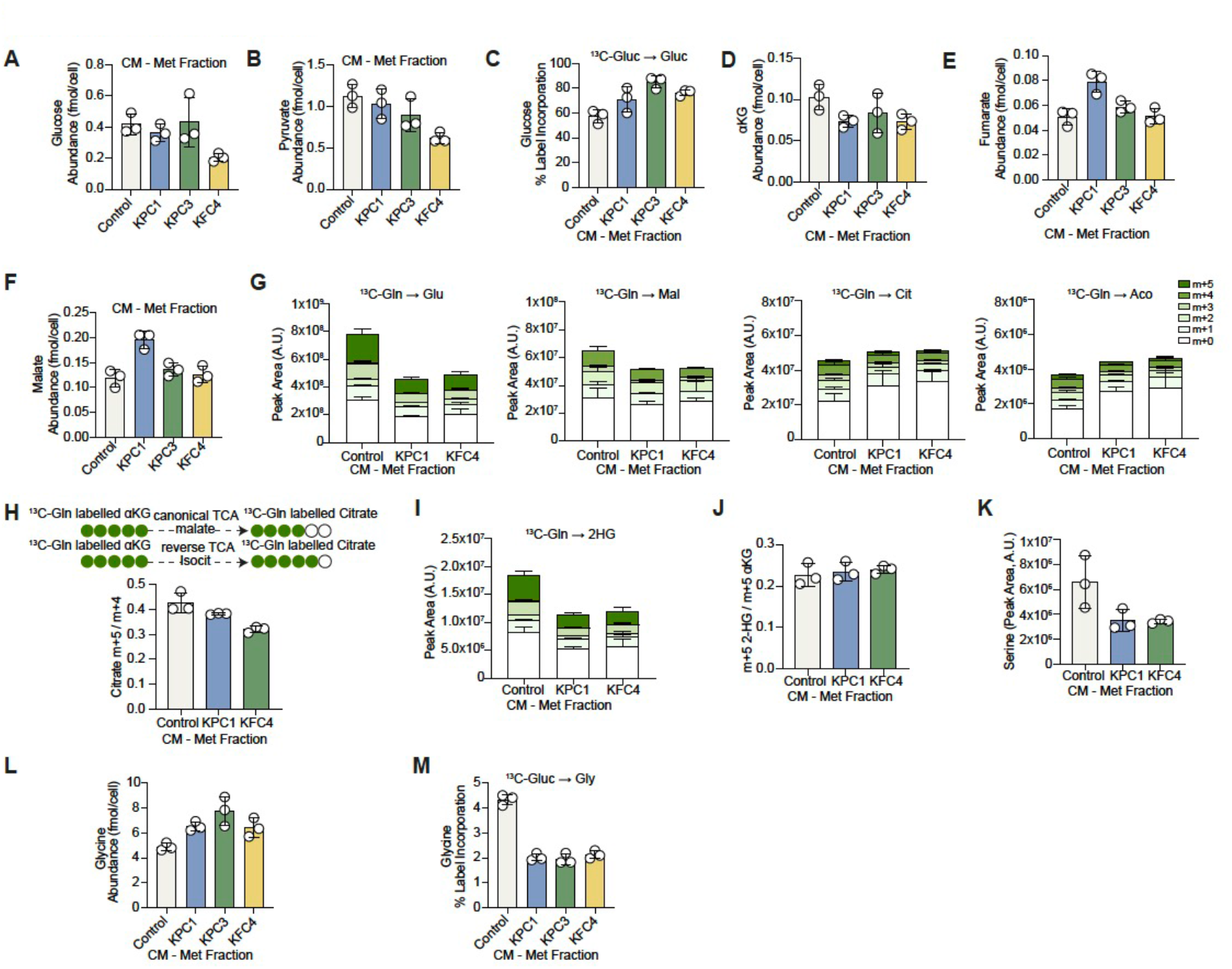
PDAC cell derived metabolites impact NAD^+^/NADH balance and decrease αKG levels in myotubes. (A) Glucose abundance (fmol/cell) in myotubes treated with control or PDAC cell derived metabolites as indicated. Values are mean±s.d. from three replicates. (B) Pyruvate abundance (fmol/cell) in myotubes treated with control or PDAC cell derived metabolites as indicated. Values are mean±s.d. from three replicates. (C) Incorporation (total label %) of [U-^13^C_6_]-D-glucose-derived, labelled carbons into glucose in myotubes treated with control or PDAC cell derived metabolites as indicated. Values are mean±s.d. from three replicates. (D) αKG abundance (fmol/cell) in myotubes treated with control or PDAC cell derived metabolites as indicated. Values are mean±s.d. from three replicates. (E) Fumarate abundance (fmol/cell) in myotubes treated with control or PDAC cell derived metabolites as indicated. Values are mean±s.d. from three replicates. (F) Malate abundance (fmol/cell) in myotubes treated with control or PDAC cell derived metabolites as indicated. Values are mean±s.d. from three replicates. (G) Incorporation of [U-^13^C_5_]-L-glutamine-derived, labelled carbons into glutamate (Glu), malate (Mal), citrate (Cit) and cis-aconitate (Aco) in myotubes treated with control or PDAC cell derived metabolites as indicated. Values are mean±s.d. from three replicates. (H) Top; simplified schematic showing glutamine derived carbon labelling pattern of citrate from canonical and reverse TCA cycle. Bottom; ratio of glutamine-derived m+5 over m+4 labelled citrate in myotubes treated with control or PDAC cell derived metabolites as indicated. Values are mean±s.d. from three replicates. (I) Incorporation of [U-^13^C_5_]-L-glutamine-derived, labelled carbons into 2-hydroxyglutarate (2HG) in myotubes treated with control or PDAC cell derived metabolites as indicated. Values are mean±s.d. from three replicates. (J) Ratio of [U-^13^C_5_]-L-glutamine-derived m+5 2-HG over m+5 labelled αKG in myotubes treated with control or PDAC cell derived metabolites as indicated. Values are mean±s.d. from three replicates. (K) Serine abundance in myotubes treated with control or PDAC cell derived metabolites as indicated. Values are mean±s.d. from three replicates. (L) Glycine abundance (fmol/cell) in myotubes treated with control or PDAC cell derived metabolites as indicated. Values are mean±s.d. from three replicates. (M) Incorporation (total label %) of [U-^13^C_6_]-D-glucose-derived, labelled carbons into glycine in myotubes treated with control or PDAC cell derived metabolites as indicated. Values are mean±s.d. from three replicates.

**Extended Data Figure 5:**
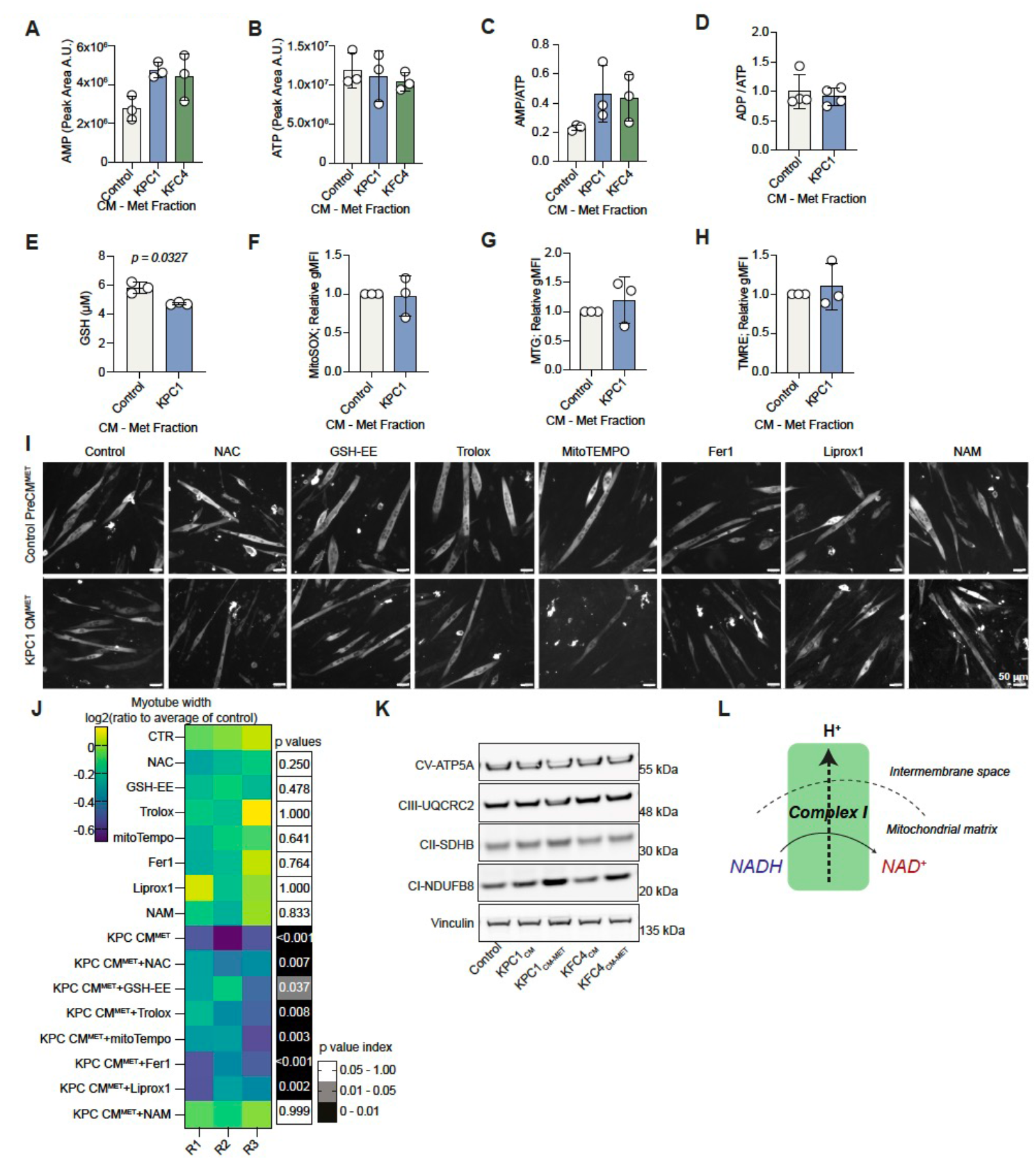
NAD^+^/NADH imbalance defines PDAC-induced myotube atrophy. (A-C) AMP **(A)**, ATP **(B)** abundance and AMP/ATP ratio **(C)** in myotubes treated with control or PDAC cell derived metabolites as indicated. Values are mean±s.d. from three replicates. (D) ADP/ATP ratio in myotubes treated with control or KPC1 cell derived metabolites. Values are mean±s.d. and average of >3 wells from four independent experiments. (E) GSH abundance (μM) in myotubes treated with control or KPC1 cell derived metabolites. Values are mean±s.d. and average of >3 wells from three independent experiments. Statistical significance was assessed with two-tailed Welch’s *t*-test. (F) Mitochondrial superoxide (MitoSOX, gMFI) levels of myotubes treated with control or KPC1 cell derived metabolites. Values are mean±s.d. and average of three wells from three independent experiments. (G) Mitochondrial content (MTG, gMFI) of myotubes treated with control or KPC1 cell derived metabolites. Values are mean±s.d. and average of three wells from three independent experiments. (H) Mitochondrial membrane potential (TMRE, gMFI) of myotubes treated with control or KPC1 cell derived metabolites. Values are mean±s.d. and average of three wells from three independent experiments. (I) Representative images of MyHC-stained myotubes treated with metabolite fraction from control or KPC1 cell conditioned media and supplemented with N-acetyl cysteine (NAC), glutathione ethyl ester (GSH-EE), Trolox, MitoTEMPO, Ferrostatin-1 (Fer-1), Liproxstatin-1 (Liprox-1) and Nicotinamide (NAM) as indicated. Scale bars; 50 μm. (J) Heatmap, myotube width quantification from **(F)** as log2 ratio to average of controls. Grayscale heatmap indicates *P* values for each condition compared to control. (K) Immunoblots of ATP5A (complex V), UQCRC2 (complex III), SDHB (complex II), NDUFB8 (complex I) and Vinculin (loading control) in myotubes treated with Control or KPC1-derived metabolites. Blots are representative of three independent experiments. (L) Scheme outlining NAD^+^ regeneration from complex I.

**Extended Data Figure 6:**
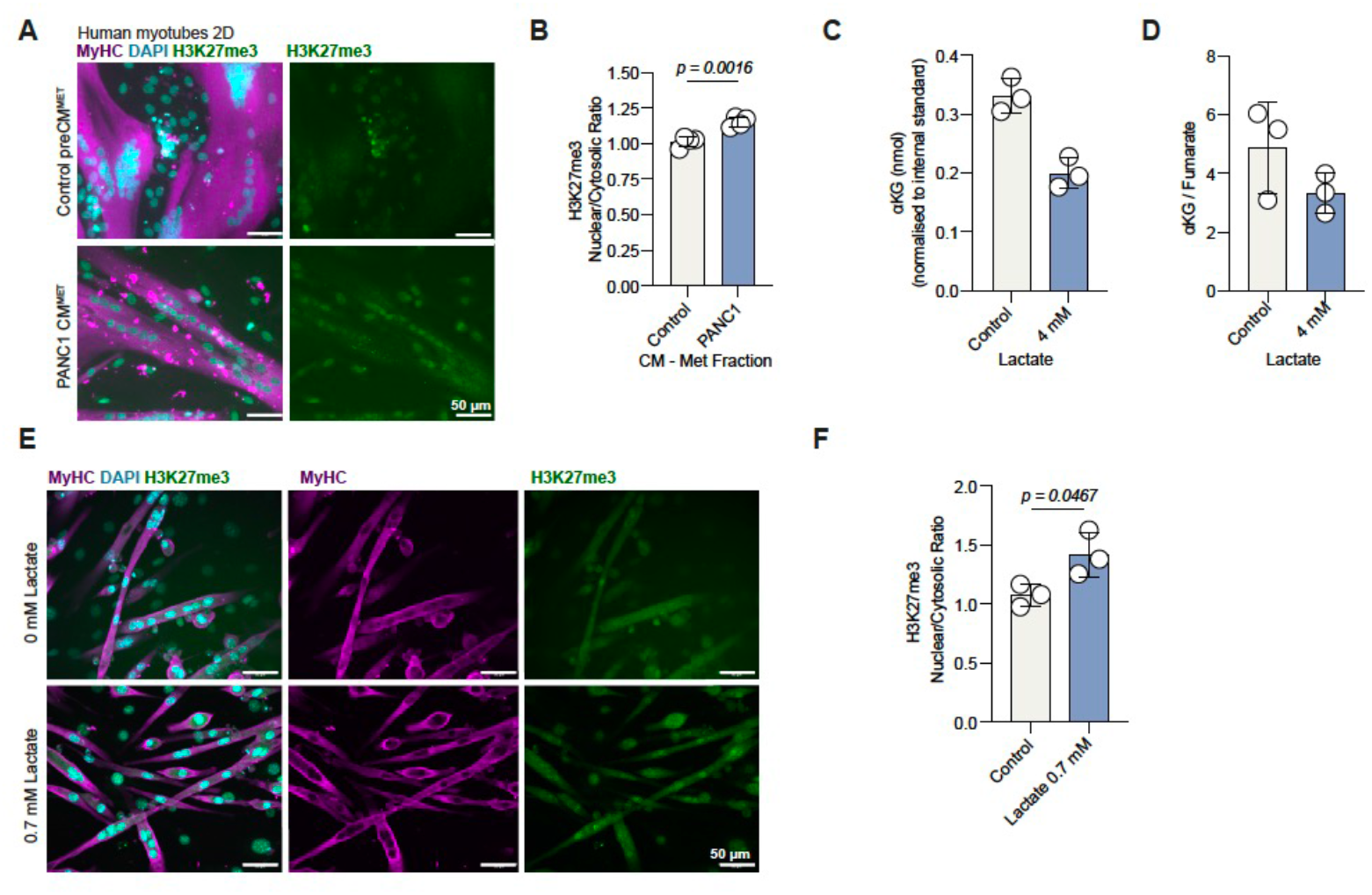
αKG levels in PDAC-induced myotube atrophy influences histone methylation. (A) Representative images of MyHC (magenta), H3K27me3 (green) and DAPI (cyan)-stained human myotubes treated with control or PANC1-derived metabolites as indicated. Scale bars; 50 μm. (B) H3K27me3 nuclear to cytosolic ratio of myotubes from **(A)**. Values are mean ± s.d. from four replicates. (C) αKG abundance (nmol) in myotubes treated with lactate as indicated. Values are mean±s.d. from three replicates. (D) αKG to fumarate ratio in myotubes treated with lactate as indicated. Values are mean±s.d. from three replicates. (E) Representative images of MyHC (magenta), H3K27me3 (green) and DAPI (cyan)-stained myotubes treated lactate (0.7 mM) as indicated. Scale bars; 50 μm. (F) H3K27me3 nuclear to cytosolic ratio of myotubes from **(E)**. Values are mean ± s.d. from three replicates. Statistical significance was assessed with two-tailed unpaired *t*-test.

**Extended Data Figure 7:**
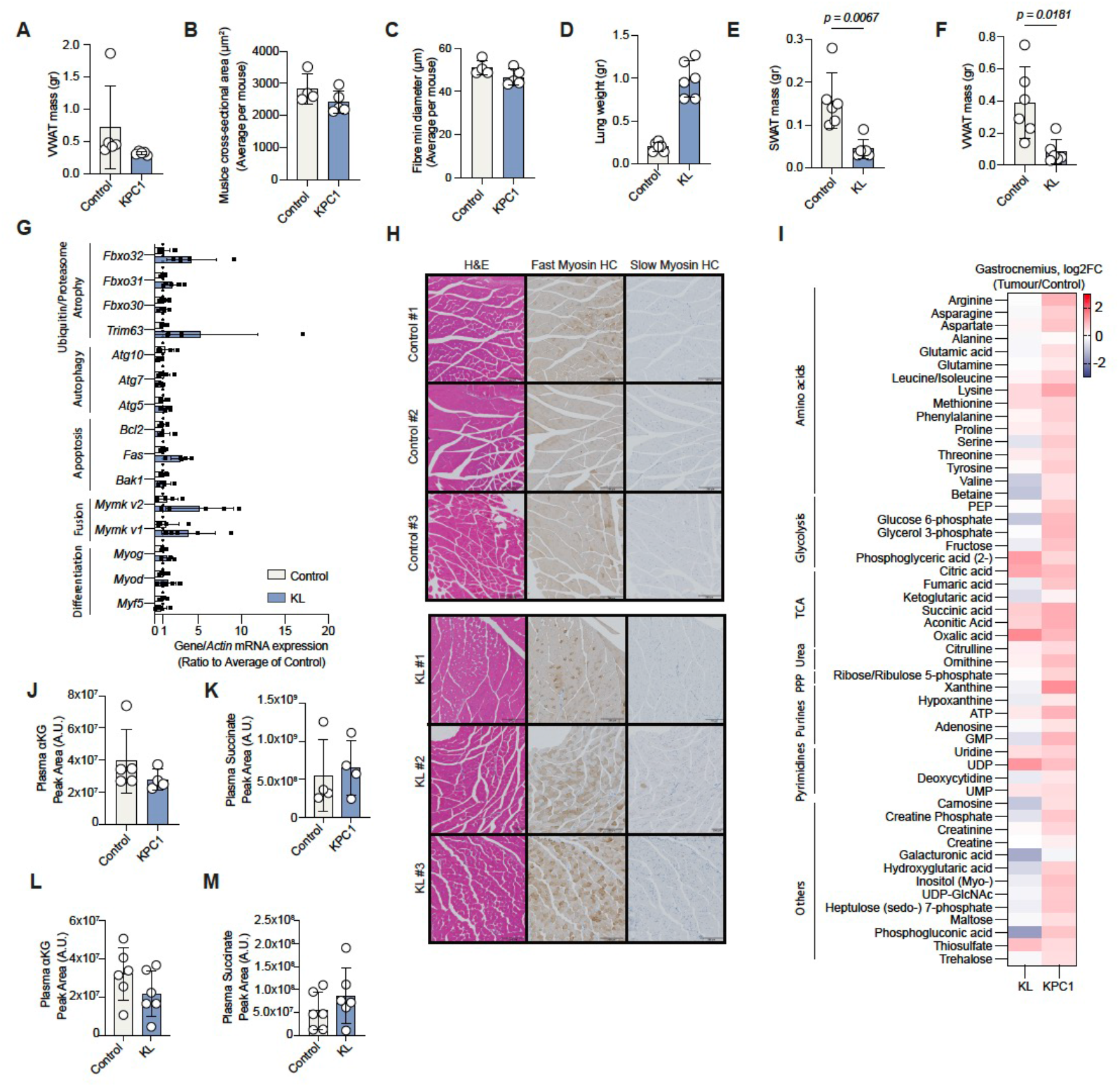
Muscle exhibits reduced αKG/succinate ratio in cancer-associated cachexia *in vivo.* (A) VWAT mass of control or KPC1 mice at endpoint. Values are mean±s.d. from n = 5 control and n = 5 KPC1 transplanted mice. (B-C) Quadriceps muscle fibre cross-sectional area (μm^2^) **(B)** and min diameter (μm) **(C)** of control or KPC1 mice at endpoint. Values are mean±s.d. from n = 4 control and n = 5 KPC1 mice. (D) Lung weight (gr) of mice from **(D)** at endpoint. Values are mean±s.d. from n = 6 control and n = 6 KL mice. (E-F) SWAT **(E)** and VWAT **(F)** mass (gr) of control or KL mice at endpoint. Values are mean±s.d. from n = 6 control and n = 6 KL mice. Statistical significance was assessed with two-tailed Welch’s *t*-test. (G) mRNA expression levels of indicated genes in gastrocnemius muscle of control or KL mice at endpoint. Values are mean±s.d. from n = 5 control and n = 5 KL mice and shown as ratio to control. Ct values were normalised to *Actin*. (H) Representative immunohistochemistry images of gastrocnemius muscle from control or KL mice showing H&E, Fast Myosin Heavy Chain and Slow Myosin Heavy Chain. Scale bars, 200 μm. (I) Heatmap, abundance of detected metabolites in gastrocnemius muscle of KL or KPC1 mice as indicated compared to their respective controls. Values are log2FC of tumour/control and are average of n = 6 control and n = 6 KL or n = 5 control and n = 5 KPC1 mice. (J) Plasma αKG abundance in control or KPC1 mice at endpoint. Values are mean±s.d. from n = 4 control and n = 4 KPC mice. (K) Plasma succinate abundance in control or KPC1 mice at endpoint. Values are mean±s.d. from n = 4 control and n = 4 KPC mice. (L) Plasma αKG abundance in control or KL mice at endpoint. Values are mean±s.d. from n = 6 control and n = 6 KPC mice. (M) Plasma succinate abundance in control or KL mice at endpoint. Values are mean±s.d. from n = 6 control and n = 6 KPC mice.

